# Global and Gene-specific Transcriptional Responses to Acute Stress

**DOI:** 10.1101/2021.07.16.452657

**Authors:** Harry Fischl, Thomas Brown, Andrew Angel, Jane Mellor

## Abstract

Nucleosomes may regulate transcription by controlling access to promoters by transcription factors and RNA polymerase II (Pol2). Potentially active genes display nucleosome depleted regions flanked by positioned -1 and +1 nucleosomes. On yeast genes, the transcription start site (TSS) is on the upstream face of the +1 nucleosome, but whether precise +1 nucleosome positioning controls Pol2 access to the TSS remains unclear. Here, using acute nutrient starvation to rapidly reprogramme the genome, we show highly dynamic upstream or downstream shifts in the position of +1 nucleosomes, coincident with levels of transcriptionally engaged Pol2 at 58% of genes. Transcript level changes broadly reflect Pol2 occupancy changes with a delay but can be further influenced by Pub1 or Puf3 dependent changes in transcript degradation rates. The response to acute stress has a second component as we also observed genome-wide changes in Pol2 distribution on genes, independent of changes in Pol2 occupancy, with Pol2 accumulating upstream of a +170 nt stalling site. Mathematical modelling supports a global increase in promoter-proximal early transcription termination as a major component of the global stress response. Thus, we uncover a two-component transcriptional response to stress, one focused on the +1 nucleosome, the second on Pol2 itself.

- A two-component responses to acute stress involving a gene-specific response and a global response
- Dynamic shifting of +1 nucleosome position with transcriptional activation or repression.
- Global targeting of Pol2 leading to early transcription termination on acute stress

## INTRODUCTION

Nucleosome positioning around the promoter is proposed to play a fundamental role in the regulation of transcription (2) by promoting or preventing access to transcription factors and RNA polymerase II (Pol2). Nucleosomes adopt well-defined positions upstream (−1 nucleosome) and downstream (+1 nucleosome) of a nucleosome-depleted region (NDR) at gene promoters (3-8). These NDRs are proposed to enable access to the DNA template at the promoter region, which is enriched for motifs recognised by sequence-specific transcription factors. The transcription start site (TSS) occurs at the 5’ boundary of the +1 nucleosome in yeast, but it is not clear whether or when the +1 nucleosome is displaced to facilitate transcription initiation or whether the position of the +1 nucleosome reflects increases or decreases in transcription. As many of the experimental approaches lack the necessary temporal and spatial resolution, it remains to be established whether this is a general mechanism of regulation or restricted to specific gene types in specific conditions (7,9). For example, instead of being correlated with transcription, nucleosome occupancy or position is often related to transcript levels (10), which may be temporally separated from the events that lead to the activation or repression of transcription, as well as being confounded by post-transcriptional regulation. Therefore, if changes are small but significant, and temporally restricted to distinct subgroups of genes, it may be impossible using genome wide data to detect these changes.

To obtain the necessary temporal resolution for this study we chose to use *Saccharomyces cerevisiae* undergoing a rapid switch from a preferred carbon source (glucose) to galactose. This induces a temporary starvation stress and slows growth, leading to a characteristic transcriptome signature known as the slow growth transcriptome signature (1,11-15) or environmental stress response (ESR) (11,16-18). These changes include a decrease in growth-related ribosomal protein gene (RPG) and ribosomal biogenesis gene (RiBi) transcripts and an increase in transcripts for genes which can be categorized as being required for stress defence, including for carbohydrate metabolism, protein folding and degradation, autophagy, and DNA damage repair (19). Cells also increase their use of aerobic respiration relative to glycolysis for ATP production with a corresponding increase in mitochondrial transcripts (20,21). While the changes to mRNA levels as a result of a transition to a slower growth rate or exposure to stress have been extensively profiled, much less is understood about Pol2 transcription during stress, the speed and nature of the transcriptional response and how this impacts the chromatin template. Once Pol2 initiates transcription, it is bound by a host of transcription elongation factors, that assist the movement of Pol2 through the chromatin template and processing of the nascent transcript (22,23) but it is not clear how nutrient stress influences the ability of Pol2 to transcribe through gene body nucleosomes and whether this is related to the position of the +1 nucleosome. Modelling and simulations of the position of Pol2 on genes as cells sense their environment can provide new insights into this question (24).

In this work we show that Pol2 occupancy on genes responds rapidly and dynamically in two distinct ways to the Glu-Gal shift: there are gene-specific responses and a global genome-wide response. The gene-specific response to stress reflects changes in promoter-associated transcription factor occupancy and the position of the +1 nucleosome, which shift relative to their position in Glu and in concordance with increased or decreased Pol2 occupancy. We were able to uncover these new associations by unbiased clustering of genes into sub-groups based on the pattern of highly dynamic increases and/or decreases in Pol2 occupancy over time, rather than dividing genes into groups based on existing gene ontology classification, or on changes to transcript levels, which show similar dynamics but with a delay and thus do not correlate as well with +1 nucleosome dynamics. Not all changes to transcript levels result simply from changes in Pol2 occupancy, as there is also a role for altered RNA stability in this response. Groups of gene transcripts that are enriched for specific RNA binding protein motifs for Pub1 or Puf3 in their 3’ UTR show some deviation from this correspondence.

The global genome-wide response to nutrient stress was uncovered by analysing base-pair resolution data on Pol2 occupancy over genes at precise time points during the Glu to Gal shift which, when combined with mathematical simulations of the shape of metagenes, reveals a shift in the promoter-proximal peak occupancy likely resulting from increased early termination of transcription. These changes may represent a general response to stress, as they occur within 5 minutes on Gal and at all genes, regardless of whether transcription, assessed by changing Pol2 occupancy decrease, increase or remain unchanged, and do not correlate with changes to the position of promoter proximal nucleosomes. Rates of transcription initiation and elongation, and/or changes to transcript stability which are linked to transcription (23,25), must change to reflect the transcriptome.

## MATERIAL AND METHODS

### Native Elongating Transcript-seq (NET-seq)

NET-seq data for Glu, Gal5, Gal15 and Gal60 were obtained from ArrayExpress E-MTAB-4568 (26). Data for two biological replicates, in which signal not specific to tagged Rpb3 had been subtracted, were obtained for each growth condition (Growth to mid-log phase (OD600 = 0.65) in YPD (1 % yeast extract, 1 % peptone, 2 % glucose) and after pelleting and subsequent resuspension in YPG (1 % yeast extract, 1 % peptone, 2 % galactose) for either 5, 15 or 60 min). NET-seq data for Gal180 for one biological replicate were obtained from Nguyen et al., 2014. In addition, two NET-seq data samples testing the effect of the pelleting and resuspension steps were generated in this study. For these, cells (BY4741 strain in which Rpb3 is FLAG-tagged) were grown in 2 × 2 L flasks to mid-log phase in 2 L YPD (1 L per flask) at 30 °C with shaking at 170 rpm before pelleting by spinning at 5000 rpm for 3 min in a JA-10 Beckman rotor, in the same way as for the experiments switching cells to YPG. Instead of resuspension in YPG, cells were either resuspended in the YPD in which they had previously been growing (sameGlu) or new YPD prewarmed to 30 °C (freshGlu) and then harvested by filtration followed by flash-freezing after a further 15 min incubation at 30 °C. For all timepoints, filtration was started prior to the timepoint to allow for the time for filtration. Cells were then flash-frozen precisely at the timepoint. NET-seq libraries were generated and sequenced as described in Fischl et al., 2017 (26).

Sequencing reads were aligned, processed and subtracted of background signal (signal that is not specific to immunoprecipitation of Pol2 via its tagged Rpb3 subunit) as described in Fischl et al., 2017 (26). As NET-seq data for both resuspension conditions were very similar to that obtained from growth in YPD, it was assumed that the background signals were also similar. Therefore, we used data obtained from ArrayExpress E-MTAB-4568, generated from carrying out the NET-seq procedure in a non-tagged BY4741 strain grown in YPD, which were used to subtract the background signal from NET-seq data from growth in YPD, to subtract the background signal in the new data. As the NET-seq data from the Gal180 sample were most similar to those obtained for Gal60, the signal used to subtract the background signal from Gal60 was used to subtract the background signal for Gal180.

### RNA-seq

Cells were grown, harvested and ground in the same way as for NET-seq. 50 mg yeast grindate was resuspended in 400 μl T.E.S. (100 mM Tris-HCl (pH 7.5), 100 mM EDTA, 0.5% SDS) and 400 μl phenol:chloroform (pH 4.7) and incubated (65 °C, 20 min, 1400 rpm). After spinning (16,000 g, 20 min, 4 °C), the upper layer was mixed with 40 μl 3 M NaOAc (pH 5.5) and 1 ml ethanol and incubated (−80 °C, 30 min). RNA precipitate was pelleted (16,000 g, 10 min, 4 °C) and resuspended in 100 μl H20. 10 μg extracted RNA was depleted of ribosomal RNA (Ribominus kit (Invitrogen)). Ion Total RNA-seq Kit v2 (ThermoFisher) was used to generate libraries for sequencing. Libraries were loaded onto the Ion Chef System (ThermoFisher) for template preparation and chip loading and the resulting chips were sequenced on the Ion Proton Sequencing System (ThermoFisher) as per the manufacturer’s instructions.

Sequencing reads were aligned to the sacCer3 genome build using the Ion Torrent Server TMAP aligner with default alignment settings (-tmap mapall stage1 map4). Non-uniquely mapped reads were removed using Samtools (samtools view -bq 1).

### MNase-seq

Cells were grown and switched to YPG as for NET-seq. To cross-link chromatin, cells were incubated for 10 min with 1 % formadehyde. Cells were then incubated for 5 min with glycine (0.125 M final). Cells were then filtered, washed with 1 % cold PBS, flash-frozen in liquid nitrogen and ground. 100 mg yeast grindate was resuspended in 500 μl MNASE buffer (20 mM HEPES pH 7.4, 0.5 % Triton-X-100, 0.1 % Tween 20, 5 mM CaCl2, complete protease inhibitors (Roche). 400 μl was incubated (15 min, 37 °C) with 8 μl MNase solution (>100 U/μl) (ThermoFisher). Reactions were stopped by adding EGTA (22.5 mM final) and incubating at 65 °C, 10 min, and cleared by spinning (16,000 g, 10 min, 4 °C). To decrosslink, the supernatant was incubated (3 h, 65 °C) with 28 μl 5 M NaCl. It was then incubated (30 min, 37 °C) with 4 μl RNase A (10 mg/ml), and (20 h, 65 °C) with 10 μl Proteinase K. DNA was purified using spin columns.

Size selection of purified DNA was carried out by electrophoresis using 8 % TBE polyacrylamide gels, selecting mononucleosomal DNA fragments (∼150-220 bp). DNA was purified from the gel band by incubation (20 h, 25 °C) in 668 μl DNA soaking buffer (0.3 M NaCl, 10 mM Tris-Cl (pH 8.0), 0.97 mM EDTA. DNA was precipitated with 750 μl isopropanol and 1 μl Glycoblue. DNA precipitate was pelleted (16,000 g, 20 min, 4 °C), washed with 80 % ethanol and resuspended in 50 μl H20. NEBNext Fast DNA library prep set for Ion Torrent was used to generate libraries for sequencing. Libraries were loaded onto the Ion Chef System (ThermoFisher) for template preparation and chip loading and the resulting chips were sequenced on the Ion Proton Sequencing System (ThermoFisher) as per the manufacturer’s instructions.

Sequencing reads were aligned to the sacCer3 genome build using the Ion Torrent Server TMAP aligner with default alignment settings (-tmap mapall stage1 map4). Bam files were further processed using DANPOS2 (27) (28) (dpos -a 1) to give genome-wide maps of the centre point of each nucleosome.

### Transcript annotations

The TSS and TES of all genes encoding mRNAs were taken from Pelechano et al., 2013 (29), selecting first for the most abundant transcript isoforms expressed in YPD and, of these, the longest, to obtain unique annotations for 5579 genes. Pol2 and RNA level gene counts were obtained by intersecting these annotations with NET-seq and RNA-seq data, respectively. Pol2 counts were rounded down to the nearest integer value and counts less than zero, as a result of non-tagged strain signal subtraction, were set to zero.

### Similarity analysis

Pol2 and RNA levels counts were normalized using the regularized log2 transformation algorithm (with blinding) within the DESeq2 R package (30). This normalizes for library size and transforms data to the log2 scale while minimizing differences between rows with low counts. Principal component analysis (PCA) on the top 500 genes with the largest variances across samples and Euclidean distance calculations were then carried out on the transformed counts.

### Differential level analysis

Differential Pol2 or RNA levels for each pairwise comparison were assessed using the DESeq algorithm within the DESeq2 R package using all available biological replicates. For each gene, this gives a value and standard error for the log2 fold change in relative levels between the two samples being compared and a p-value, adjusted for multiple testing using the Benjamini-Hochberg method (padj), showing the significance of any difference, or two-fold difference, depending on the threshold that was set.

### Pol2 level profile clustering

Regularized log2-transformed Pol2 level counts (without blinding) for Glu, Gal5, Gal15 and Gal60 samples were averaged for each pair of biological replicates. To enable comparison between genes, these counts were then standardized across the time series to give a mean and standard deviation of 0 and 1, respectively. Genes showing a significant two-fold change in Pol2 level between a point in the time series and any subsequent point were then clustered into 5 groups using the fuzzy c-means soft clustering algorithm implemented by the Mfuzz R package (31,32). Cluster membership values, which indicate how well a profile matches a cluster, were used to assign each gene profile to a cluster and exclude gene profiles without a membership value greater than 0.6 for any cluster.

### Motif analysis

Position Weight Matrices (PWMs) for the sequences recognized by selected *S. cerevisiae* transcription factors were taken from the JASPAR database http://jaspar.genereg.net/. Promoter sequences (TSS-500 bp to TSS+99 bp) were scanned for matches to these sequences using a significance threshold of <0.001. Both strands were scanned for all motifs apart from Sfp1, which has been shown to display directionality (33). Overlapping motifs on either strand were combined and counted as 1 motif with the central position recorded as its location. The number of motifs per distal (TSS-201 bp to TSS-400 bp) and proximal (TSS-1 bp to TSS-200 bp) promoter region was then counted for all genes from each cluster. Significant (p value < 0.05 after adjusting for multiple testing using the Holm method) enrichment or depletion of the motif in genes of a cluster relative to all other genes not in any cluster was determined using a Poisson rate test.

### Transcription factor analysis

Data for Msn2 ChIP-seq from cells grown in media containing 2 % glucose and 20 min after switching to glucose-depleted media were taken from Elfving et al., 2014 (34). Paired end fastq files were aligned using HISAT2 (hisat2 --no-spliced-alignment --no-mixed --no-discordant) to the sacCer3 genome. Bam files of aligned reads were then converted to reads per million normalized bigwig coverage files. The log2 fold change in Msn2 levels were then calculated for each gene for 50 bp bins across the full promoter window (TSS-500 bp to TSS+99 bp) and the mean fold change for each bin across all genes within each cluster was then calculated after first removing the top and bottom 5 % of values. The log2 fold change was also calculated for the proximal promoter window (TSS-200 bp to TSS-1 bp) and a significant (p value < 0.05 after adjusting for multiple testing using the Holm method) difference in the mean fold change for all genes within a cluster relative to all other genes not in any cluster was determined using a Welch’s two-tailed t-test after first removing the top and bottom 5 % of finite values for each grouping.

Data for RNA levels at multiple time points during a 1 h time period following exposure to H2O2 for both wildtype and *msn2msn4* strains were taken from Huebert et al., 2012 (35). The average RNA level for each cluster was calculated for each time point and for each strain. Significant differences in the mean RNA level between wild type and *msn2msn4* strains and between clusters 3 and 4 were determined at the 12 min timepoint using a Welch’s two-tailed t-test.

Data for RNA levels at multiple time points during a 1.5 h time period following specific induction of Msn2 expression were taken from Elfving et al., 2014 (34). The average RNA level for each cluster was calculated for each time point. Significant increases in the mean RNA level for all genes within a cluster relative to all other genes not in any cluster and for cluster 4 relative to cluster 3 were determined using a Welch’s one-tailed t-test.

Data for Sfp1 ChEC-seq were taken from Alberts et al., 2019 (33) using the data in which the activity of MNase fused to Sfp1 was induced for 240 s and a control in which MNase activity was induced for 20 min but was not fused to Sfp1. The log2 fold change in Sfp1 signal relative to the control was calculated for each gene for 50 bp bins across the full promoter window (TSS-500 bp to TSS+99 bp) and the mean fold change for each bin across all genes within each cluster was then calculated after first removing the top and bottom 5 % of values. The log2 fold change was also calculated for the proximal (TSS-200 bp to TSS-1 bp) and distal (TSS-400 bp to TSS-201 bp) promoter windows. A significant (p value < 0.05 after adjusting for multiple testing using the Holm method) difference in the mean fold change for all genes within a cluster relative to all other genes not in any cluster was determined using a Welch’s two-tailed t-test after first removing the top and bottom 5 % of finite values for each grouping.

Data for RPM normalized Rpb1 ChIP-seq bedgraph tracks before and after over expression of Sfp1 for 1 h and before and after anchoring away Sfp1 for 1 h were taken from Alberts et al., 2019 (33). The log2 fold change in Rpb1 level was determined for each gene for the window from TSS to TES. A significant (p value < 0.05 after adjusting for multiple testing using the Holm method) difference in the mean fold change for all genes within a cluster relative to all other genes not in any cluster was determined using a Welch’s two-tailed t-test after first removing the top and bottom 5 % of finite values for each grouping.

### Nucleosome shift analysis

The position of the centre point of the +1 nucleosome for each gene was determined from the DANPOS2 output as the centre point of the nucleosome located within the window TSS to TSS+100 bp (28). For genes with two centre points within this window, the one closest to TSS+50 bp was used. The difference between this position between Glu and each Gal time point (Gal5/15/60) was then determined. Significant (p value < 0.05 after adjusting for multiple testing using the Holm method) mean differences in +1 nucleosome positions were determined for each of the three conditions for genes within each cluster using a one sample two-tailed, t-test. Genes in which there was no nucleosome centre point located within the TSS to TSS+100 bp window in either the Glu or Gal condition being analysed were excluded.

### Elongation profile analysis

For heatmaps of Pol2 levels over genes, for each gene, the mean NET-seq counts in 10 bp bins from the TSS-100 bp to the TES+100 bp were calculated. Mean counts less than zero were set to zero. To normalize for differences in total Pol2 levels over genes, mean counts for each bin were standardized by subtracting the mean count for bins within the window TSS to TSS+1000bp and dividing by the standard deviation for bins within this window.

Metagenes showing the average profile across genes for each condition were calculated as follows. For metagenes of the region from TSS to TSS+1000 bp (early gene profile), counts within the window TES to TES-200 bp were excluded as were genes shorter than 200 bp. For metagenes of the region from TES-300 bp to TES (late gene profile), counts within the window TTS to TTS+500 bp were excluded as were genes shorter than 600 bp. These exclusions prevent the profiles near the TES and TTS influencing the metagenes of early and late gene profiles, respectively. The mean NET-seq counts in 10 bp bins from the TSS-100 bp to the TES+100 bp were calculated for the early gene profile and from the TES+100 bp to the TSS-100 bp for the late gene profile. Mean counts less than zero were set to zero. Both bins for the early and late gene profiles standardized by subtracting the mean count for bins within the window TSS to TSS+1000bp and dividing by the standard deviation for bins within this window. Mean counts were then calculated for each bin across all genes or subset of genes after first removing the top and bottom 5 % of bins.

For travelling ratio calculations genes shorter than 701 bp were excluded. For each gene the mean count for the 25 10 bp bins within the window from TSS to TSS+250 bp was divided by the mean count for the 25 10 bp bins within the window from TSS+251 bp to TSS+500 bp. Significant (p < 1×10^−10^) differences in the mean log2 transformed travelling ratio for each Gal condition (Gal5/15/60) relative to Glu were determined using a Welch’s two-tailed t-test.

Heatmaps, metagenes, and travelling ratios were calculated on NET-seq data produced from combining the 2 biological replicates for each condition. Metagenes and travelling ratios were also calculated from individual biological replicates.

### Degradation rate

Regularized log2-transformed Pol2 level counts (without blinding) for Glu and Gal60 samples were averaged for each pair of biological replicates. Regularized log2-transformed RNA level counts (without blinding) for Glu and Gal60 samples were averaged for each pair of biological replicates. These transformed RNA level counts were then subtracted from these transformed Pol2 level counts to give the log2-transformed ratio of Pol2 to RNA level for both Glu and Gal60 conditions that we have used as the relative log2-transformed degradation rate for each condition.

### Gene ontology analysis

Gene ontology (GO) analysis was performed using the gost algorithm within the gprofiler2 R package. The 5579 genes analysed in this study were used as a custom background gene list for analysis of functional enrichment in each of the NET-seq clusters. For GO analysis on genes belonging to each NET-seq cluster and also showing increased mRNA stabilisation in either glucose or galactose as well as a significant difference in RNA level at least at one time point during the hour following the switch to YPG, a custom background gene list of these genes showing a significant difference in RNA level was used.

### Motif enrichment

Motif enrichment analysis was performed using DREME (36). For promoter motif enrichment analysis, Fasta files containing the set of sequences for the window TSS-500 bp to TSS-1 bp for all genes from a cluster were compared to sequences from other clusters as explained in the results. The reverse complement of the sequences were also searched. For 3’UTR motif enrichment analysis, Fasta files of the sequence from the region from the end of the coding sequence to the TES were used. Genes with 3’UTRs less than 50 nucleotides were excluded. Genes showing increased stability in Glu were compared against those showing increased stability in Gal60 and vice versa. Genes were also required to show a significant difference in RNA level at least at one time point during the hour following the switch to YPG.

### Mathematical modelling

The process of transcription was formulated as a stochastic process with core components of initiation; elongation; polymerase occlusion; stalling; resumption of elongation from the stalled state; backtracking from a stalled state; resumption of elongation from the backtracked state; collision-induced stalling and termination; early termination with a Poisson distribution around a fixed location; two dynamic windows, in which there can be different stalling, backtracking and resumption rates; termination at the 3’ end of a gene. When polymerases collide, the situation resolves itself depending on the state of the polymerases involved. If a moving polymerase collides with another moving polymerase, the upstream polymerase becomes stalled. If a moving polymerase collides with a stalled polymerase, the upstream polymerase will become stalled and the downstream stalled polymerase will be terminated. If a moving polymerase collides with a backtracked polymerase, the upstream polymerase will become stalled.

The simulations were limited to the beginning of a synthetic gene, which covered 1000 nt. Polymerases had a fixed footprint of 40 nt. Upon reaching the end of the synthetic gene and elongating, polymerases were removed. 150,000 parameter sets were sampled uniformly between the maximum and minimum parameter values given in Table 1, via latin-hypercube sampling (37) and each of these was simulated for a population of 100,000 identical synthetic genes. Simulations were run for the equivalent of 40 minutes in increments of 0.005 minutes per time-step to allow the system to reach a steady state. The output distribution of transcriptionally-engaged polymerase for a given parameter set was then taken of the sum of the locations of polymerases at the final time-step of each of the 100,000 simulated genes.

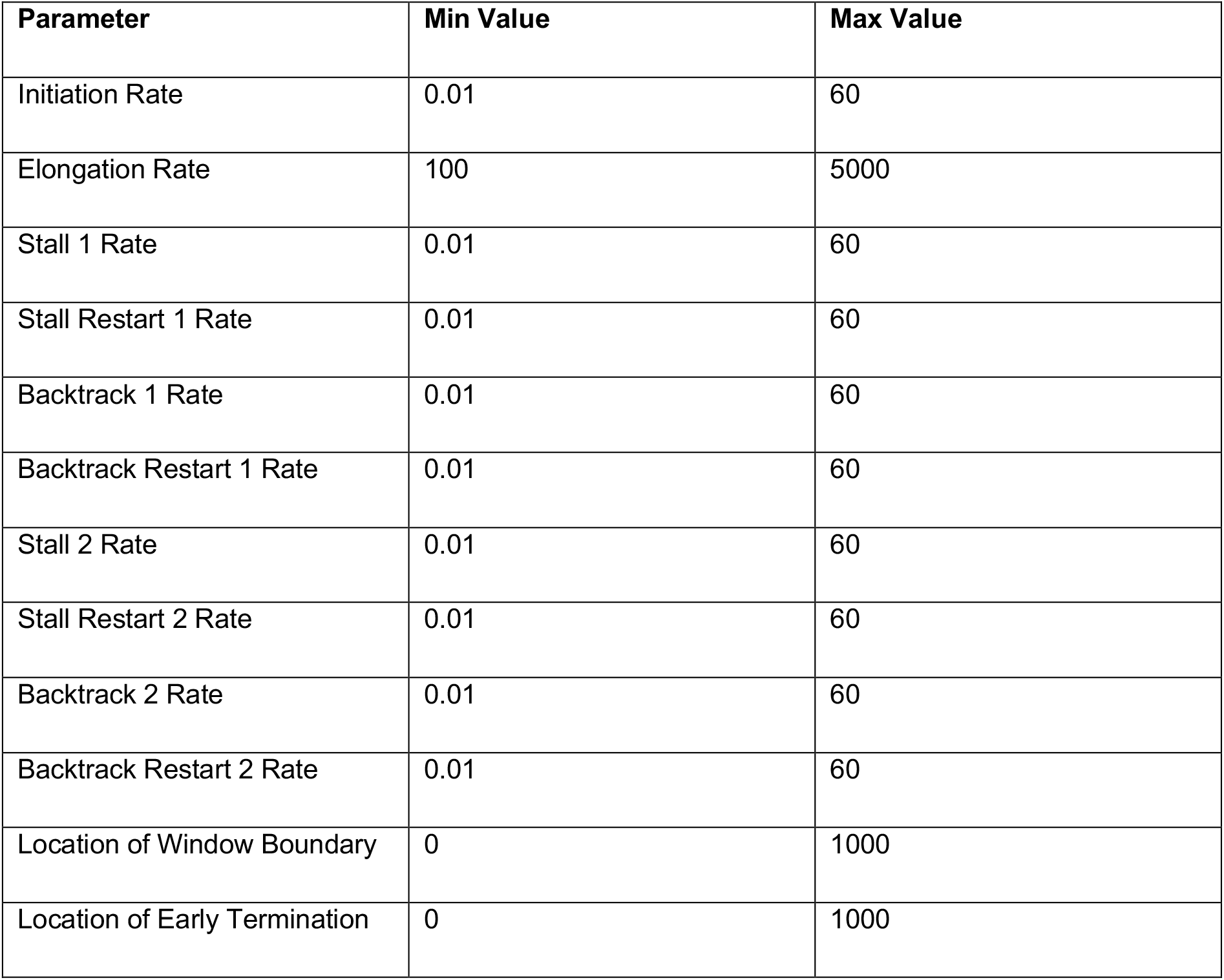

For the purposes of fitting, simulation and experimental data were binned, with bins of size 10 nt. For the experimental NET-seq data, genes were defined via annotations derived from TIF-seq (29): the TSS and TTS for each gene was defined by choosing the most abundant start and end point detected in YPD. Only genes longer than 1000 nt in length were selected and, of those, only ones with total read counts in the first 1000 nt greater than the average for all initially selected genes. Experimental and simulated NET-seq data were normalised by dividing each bin by the total read counts or sampled polymerase locations in the first 1000 nt, respectively.

Simulated data was compared to NET-seq data using the Kolmogorov-Smirnov statistic (maximum of the differences between individual points of the CDF of each data set as the goodness-of-fit metric. For the plots in Figure S6E,F, the single best fitting simulation for each gene was used; for the parameter comparisons, the 100 best fitting simulations for each gene were used.

### Sources of codes for statistical and data analysis

|  |  |  |
| --- | --- | --- |
| Galaxy Web-based platform | Usegalaxy.org | RRID:SCR_006281 |
| FastQC | http://www.bioinformatics.babraham.ac.uk/projects/fastqc/ | RRID:SCR_014583 |
| Bowtie for Illumina | (38) | RRID:SCR_005476 |
| R Project for Statistical Computing | (39) | https://www.r-project.org/;<br>RRID:SCR_001905 |
| Bioconductor | R studio | RRID:SCR_006442 |
| GenomicRanges | (40) | RRID:SCR_000025 |
| GenomicFeatures | Bioconductor | RRID:SCR_016960 |
| DEseq2 | (41) | RRID:SCR_015687 |
| DANPOS2 | (30) | RRID:SCR_015527 |
| MATLAB | (27,28) The MathWorks Inc. | RRID:SCR_001622 |
| Position Weight Matrices (PWMs) for the sequences recognized by selected *S. cerevisiae* transcription factors | http://jaspar.genereg.net/. | RRID:SCR_003030 |

## RESULTS

### Transcriptome changes strongly correlate with the slow growth signature after 15 min of a carbon source shift

*S. cerevisiae* cells grown in typical laboratory, asynchronous, batch culture conditions of aerated, glucose-containing rich media (YPD, yeast extract, peptone, 2 % glucose) are able to obtain sufficient nutrients to support a phase of exponential growth.

Due to the presence of glucose and absence of galactose, these cells do not express the genes necessary to convert galactose to glucose-6-phosphate for use in glycolysis. Sudden replacement of YPD for galactose-containing media (YPG) stresses the cells as they no longer have access to a carbon source that can be used to support the same growth rate (18). Indeed, transcription of genes required for Gal metabolism show only minimal induction, even after 60 min, and GFP-tagged Gal10 protein only appears between 4 and 6 hours after induction, demonstrating that nutrient accessibility is severely restricted for several hours following this carbon source shift (Fig. S1). To establish the timing with which other transcriptome changes occur in response to this carbon source shift, we used RNA-seq to monitor RNA levels genome-wide in cells during exponential growth in YPD (Glu) and after 5 min (Gal5), 15 min (Gal15) and 60 min (Gal60) in YPG (Gal). Comparison of the RNA levels for protein coding transcripts across all these samples using Euclidean distance and PCA shows RNA levels after 5 min are more similar to those in Glu than after 15 and 60 min (Fig. 1A,B). The slow growth transcriptome signature (1) is proposed to be tightly linked to the stress response (11). As a result, by comparing it to the transcriptome changes, we used it to gauge the degree to which the transcriptome has responded to the carbon source shift and stress at each time point (Fig. 1C). There is a positive correlation between the slow growth signature and the changes in RNA levels for all Gal samples relative to Glu (Spearman correlation coefficients ρ = 0.49 (Gal5), 0.72 (Gal15), 0.72 (Gal60)), showing that the transcriptome has begun to respond after 5 min, showing a greater response after 15 min. The positive correlation between the changes in RNA levels from Gal5 to Gal15 with the slow growth signature (ρ = 0.65) further highlights that only minor changes to the stress-dependent component of the transcriptome have begun to occur after 5 min relative to those occurring after 15 min. RNA level changes between Gal15 and Gal60 show no correlation with the slow growth signature (ρ = 0.03), suggesting the transcriptome has reached an approximate steady state by 15 min. Further confirming these observations, the number of genes showing significant differential RNA levels is low (494 genes) after 5 min, while over 2690 genes show differential RNA levels after 15 and 60 min (Fig. 1D).

**Figure 1.**
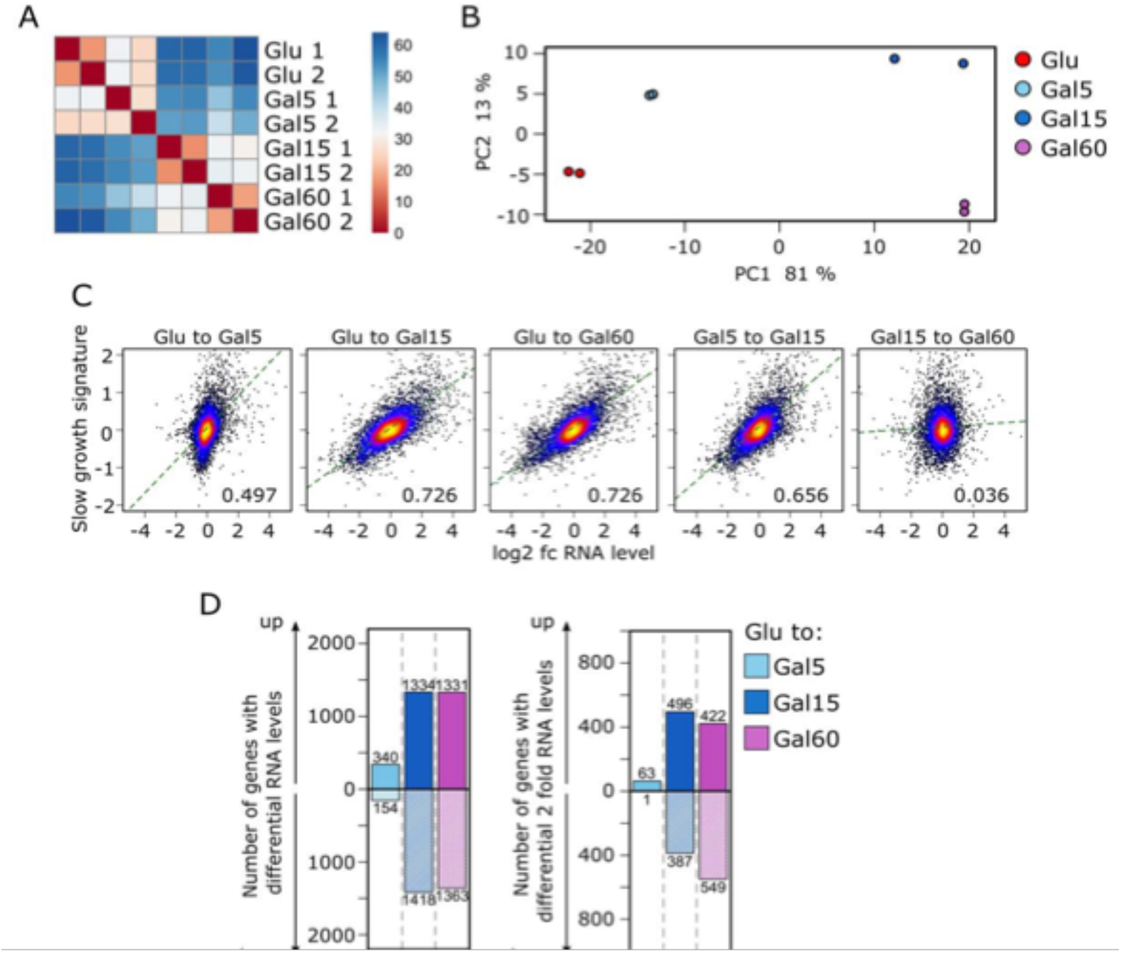
Transcriptome changes strongly correlate with the slow growth signature after 15 min of a carbon source shift. A and B) Similarity between regularized log transformed RNA level gene counts of RNA-seq samples from cells grown exponentially in YPD (Glu) and after 5 (Gal5), 15 (Gal15) and 60 min (Gal60) following the replacement of YPD for YPG (2 biological replicates for each condition) measured by the Euclidean distance between samples (A) and principal component analysis (B). C) Relationship between the slow growth signature from O’Duibhir et al., 2014 (1) and the log_2_ fold change in mRNA level for the comparisons indicated. Points are color-coded from low to high density (black < blue < red < yellow). Regression line (green) and Pearson correlation coefficient are shown. D) Number of genes showing significant (adjusted p < 0.05, Wald test) differential (left panel) or two-fold differential (right panel) RNA levels following the transfer of cells from YPD to YPG for 5 (Gal5), 15 (Gal15) and 60 min (Gal60). See also Figure S1.

### Rapid changes in Pol2 levels in response to a carbon source shift

Having established the timing with which the transcriptome changes in response to this carbon source shift and shown that it is established by 15 minutes, we next aimed to assess how these relate to changes in transcription, assessed using NET-Seq, which maps the position of transcriptionally engaged Pol2 at single nucleotide resolution in native conditions from cells that have been harvested by filtration and flash frozen in liquid nitrogen (18,22,42). As cells are harvested from native conditions without the need for labelling windows or exposure to cross-linking agents, the technique affords the ability to profile transcription levels at short precise time points. We monitored Pol2 levels genome-wide in cells during exponential growth in YPD (Glu) and after 5 min (Gal5), 15 min (Gal15), 60 min (Gal60) and 180 min (Gal180) following a switch to YPG (Gal) (18,22) (Fig. 2A-C).

**Figure 2.**
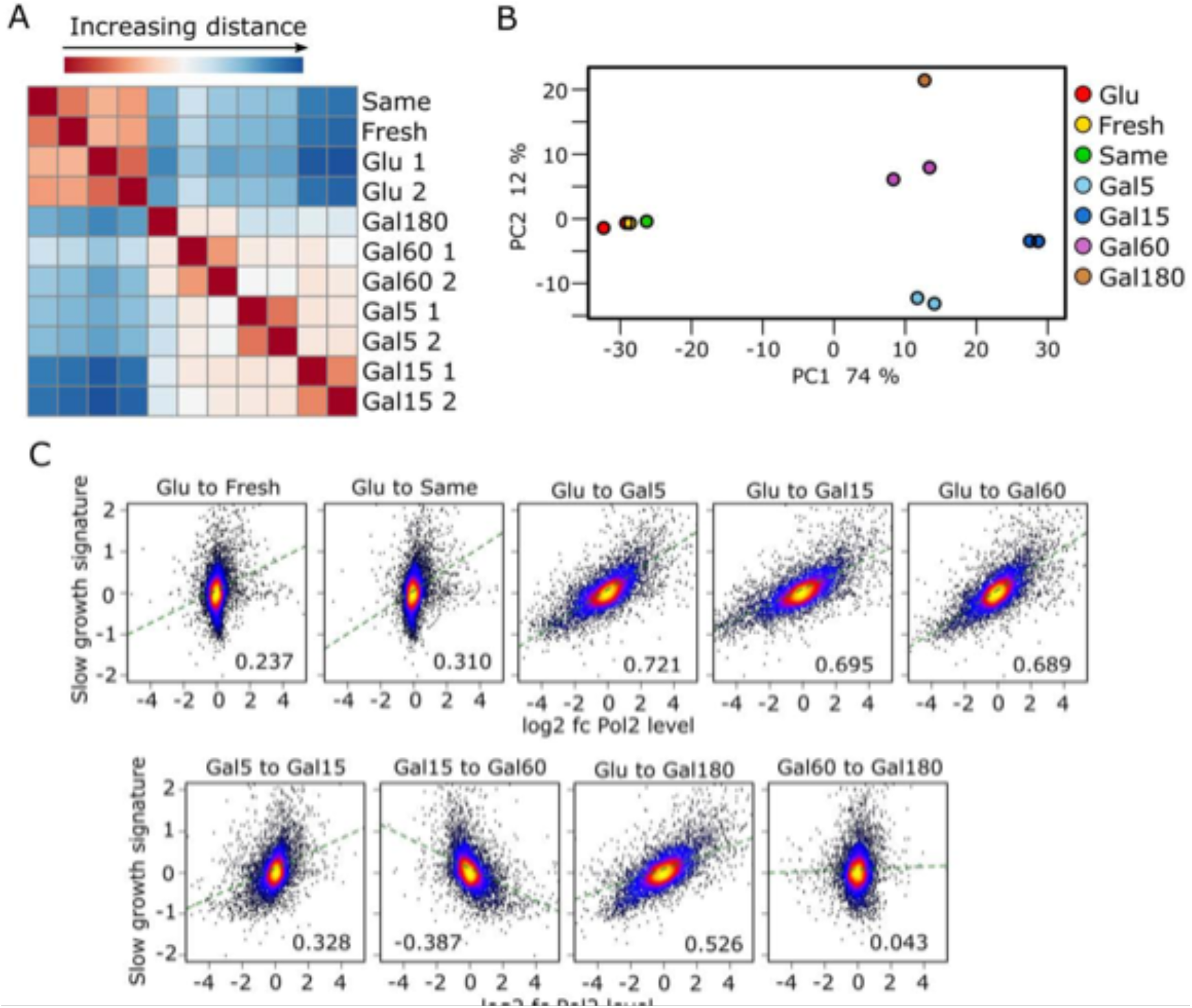
Rapid changes in Pol2 levels in response to a carbon source shift. A and B) Similarity between regularized log2 transformed Pol2 level gene counts of NET-seq samples from cells grown exponentially in YPD (Glu) and after 5 (Gal5), 15 (Gal15) and 60 min (Gal60) following the replacement of YPD for YPG (2 biological replicates for each condition) measured by the Euclidean distance between samples (A) and principal component analysis (B). C) Relationship between the slow growth signature from O’Duibhir et al., 2014 (1) and the log2 fold change in Pol2 level at mRNA genes for the comparisons indicated. Points are color-coded from low to high density (black < blue < red < yellow). Regression line (green) and Pearson correlation coefficient are shown. D) Number of genes showing significant (adjusted p < 0.05, Wald test) differential (left panel) or two-fold differential (right panel) Pol2 levels following the transfer of cells from YPD to YPG for 5 (Gal5), 15 (Gal15) and 60 min (Gal60).

In order to switch the growth medium, cells were pelleted by spinning for 3 min at 5000 rpm before resuspension in pre-warmed YPG. To control for the effect of these pelleting and resuspension steps, NET-seq data were also obtained from cells either resuspended, after spinning, in the original YPD in which they had previously been growing (sameGlu) or new pre-warmed YPD (freshGlu) and then harvested after 15 min. For these controls, changes in Pol2 levels over protein coding genes show a weak positive correlation with the slow growth signature, indicating that both cause a mild stress response (ρ = 0.31, 0.23) (Fig. 2C). Given that such pelleting and resuspension steps are typical of many laboratory protocols, it is important to note that these perturbations stress the cells to an extent that influences Pol2 occupancy on genes, possibly as a result of hypoxia (43).

Next, we compared the Pol2 levels over protein coding genes following the switch to galactose. Euclidean distance and PCA show a clear distinction between samples harvested from glucose and all those harvested following the switch to galactose (Fig. 2A-B). In addition, we examined the changes in Pol2 levels over genes with the slow growth signature (Fig. 2C). There is a strong correlation with this signature for the Pol2 level changes observed in cells at all time points following transfer from YPD (Glu) to YPG (Gal) (ρ = 0.72 (Gal5), 0.69 (Gal15), 0.68 (Gal60)). Furthermore, as changes from Gal5 to Gal15 and from Gal15 to Gal60 show positive (ρ = 0.32) and negative correlations (ρ = - 0.38), respectively, with this signature, the response to the stress at the transcriptional level can be seen to be greatest at the 15 min timepoint. This is also apparent by PCA, in which the Gal15 samples are furthest from the Glu samples along principal component 1 which explains 74 % of the variance between samples (Fig. 2B). By 60 min, genes repressed and induced in response to this carbon source shift are, in general, no longer repressed and induced, respectively, at the transcriptional level, to the same extent as after 15 min. Interestingly, this trend does not continue, as Pol2 level changes from Gal60 to Gal180 show no correlation with the slow growth signature (ρ = 0.04). This suggests that the transcription levels at most genes associated with the response to this carbon source shift have reached an approximate steady state across this 2h period. This is despite the large increase in the transcription of genes for galactose metabolism (Gal1, 7 and 10) during this window (Fig. S1B), which might be predicted to stimulate a transcription profile with greater similarity to that observed for exponentially growing cells in YPD. However, we note that Gal10 protein levels are only just becoming induced at this time (Fig. S1D,E).

Analysis of the number of genes showing significant differential Pol2 levels upon the switch to YPG emphasises the speed, scale and degree to which transcription changes (Fig. 2D). Within 5 min, 3284 genes (58 % of genes) show significantly increased or decreased levels. By 15 min even more genes show Pol2 level changes (3591 genes), a difference which is particularly apparent if restricted to significant two-fold changes (Glu to Gal5: 1130 genes; Glu to Gal15: 1578 genes). By 60 min, there is a reduction in the number of genes showing significant changes (Glu to Gal60: 2887), and two-fold changes (Glu to Gal60: 866).

### Altered transcript degradation rates fine tune transcript levels in response to stress

To explore the relationship between transcription and transcript levels in greater detail, we compared changes in Pol2 levels against changes in RNA levels for each of these 3 Gal timepoints relative to Glu (Fig. 3A). By assessing the gradients of the regression lines, large changes in Pol2 levels have only a small impact on RNA levels after 5 min. The gradient of this line becomes progressively steeper over the hour and is closest to a 1 to 1 relationship after 60 min. In order to promote rapid readjustment of the transcriptome, transcriptional changes observed at 15 min may exceed those necessary to maintain the stress-dependent component of the transcriptome at the appropriate level. Once this adjustment has taken place, Pol2 levels may then recover to those observed from 60 min onwards. Thus, while RNA level changes broadly reflect Pol2 level changes with a delay, suggesting that changes in RNA synthesis rates are the major determinant of the observed transcriptome changes, we also wanted to determine the role played by changes in RNA degradation rates.

**Figure 3.**
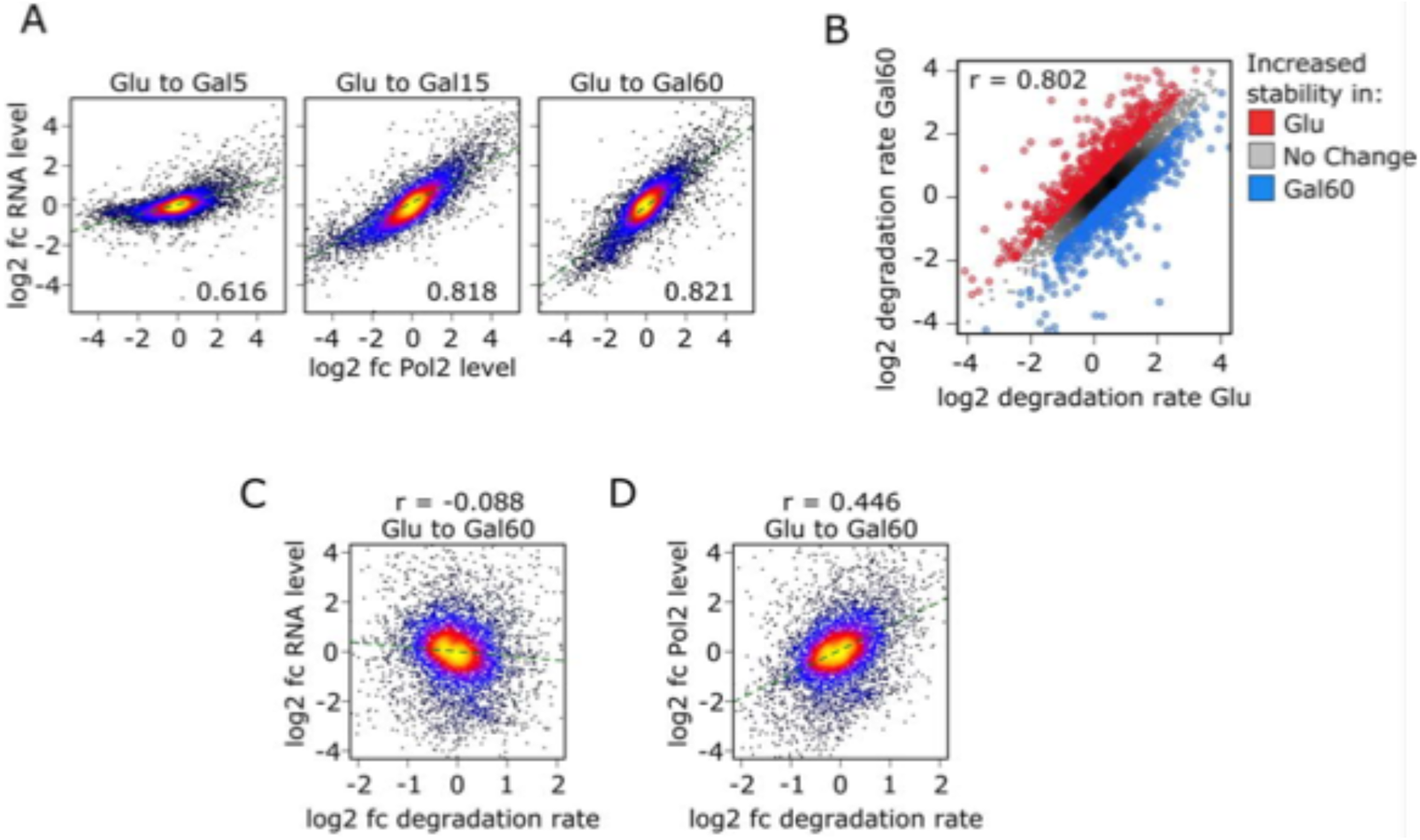
Transcript degradation rates correlation with Pol2 but not transcript levels. A) Relationship between the log2 fold change in RNA level and Pol2 level at mRNA genes for the comparisons indicated. Points are color-coded from low to high density (black < blue < red < yellow). Regression line (green) and Pearson correlation coefficient are shown. B) Relationship between the log2 transformed mRNA degradation rate in Gal60 and in Glu for all genes showing a significant change in RNA level between any of the Glu, Gal5, Gal15 and Gal60 conditions. Points are shaded from low to high density (gray < black). mRNAs showing an increase or decrease in log2 transformed degradation rate from Glu to Gal60 of at least 0.5 are highlighted in red or blue, respectively. Pearson correlation coefficient is shown. Relative log2 transformed degradation rates are calculated by subtracting the mean regularized log2 transformed RNA level from the mean regularized log2 transformed Pol2 level for each condition. C and D) Relationship between the log2 fold change in RNA level (C) or Pol2 level (D) and the log2 fold change in degradation rate from Glu to Gal60 for all genes. Points are color-coded from low to high density (black < blue < red < yellow). Regression line (green) and Pearson correlation coefficient are shown.

In steady state conditions, the mRNA degradation rate is equal to the mRNA synthesis rate divided by the total mRNA level. Under the assumption that the Pol2 levels over the gene determined by NET-seq correspond to synthesis rates, the degradation rate is then proportional to the ratio between NET-seq and RNA-seq signals.

During exponential growth, the synthesis rates and total mRNA levels are unchanged relative to the number of cells within the population and so each can be viewed as being at a steady state. Our data suggest that an approximate steady state for both Pol2 levels and RNA levels has been reached after 60 min in galactose. We were therefore able to calculate relative mRNA degradation rates for cells in Glu and Gal60. Comparing these rates shows that they strongly correlate (ρ = 0.80) between the two conditions (Fig. 3B). This shows that generally degradation rates do not show large changes in response to this change in conditions. This provides further evidence that changes in transcription are the main determinant of the change in mRNA levels in response to this stress rather than changes in mRNA stability, in disagreement with previous reports (44). There is however some modulation of mRNA degradation rate. To understand how changes in degradation rate affect gene expression, we correlated the log_2_ fold change in degradation rate with that of the RNA level (Fig. 3C). While on average an increase in degradation rate corresponds to a decrease in RNA level, this is a surprisingly weak association (ρ = 0.08). Correlating the log_2_ fold change in degradation rate with that of the Pol2 level shows a stronger correlation (ρ = 0.44) (Fig. 3D). As increased degradation rates correspond with increased synthesis rates (23,25), this may explain why increased degradation rates do not lead to greater decreases in RNA level but do reflect Pol2 occupancy.

### Genes can be clustered into functional groups with distinct patterns of induction and repression

The precise nature by which NET-seq maps transcriptional changes gives a unique insight into the patterns of induction and repression of transcription during the response to a particular stress.

We took the 1809 genes showing at least a significant two-fold change in Pol2 levels between one point and at least one other subsequent point in the time series (Glu, Gal5/15/60). Using a fuzzy c-means clustering algorithm (32), 83 % of these genes (1494 genes) could be grouped into 5 clusters according to their profile across this time series of their standardised log_2_ normalized Pol2 levels (Fig. 4A; S4A-C). These can be categorised as genes that are either slow (Cluster 1, 403 genes) or fast (Cluster 2, 330 genes) to reach their minimal levels or genes that are either slow (Cluster 3, 340 genes) or fast (Cluster 4, 348 genes) to reach their maximal levels. Genes in Clusters 2 and 4 also show recovery to a greater extent than Clusters 1 and 3 to their levels of transcription shown before stress after 60 min. Finally, there is a smaller group of genes (Cluster 5, 73 genes) that shows maximal induction by 5 min before showing repression back to close to its level in Glucose by 15 min. From 60 min to 180 min, the level of Pol2 over genes remains broadly the same for all clusters providing further evidence that the cells have reached an approximate steady state during this period (Fig. S4A). We applied each of the 5 clusters to gene ontology analysis to assess whether genes with specific functions show particular patterns of induction and repression (Table S1). Both Clusters 1 and 2 are enriched for RiBis, while RPGs are enriched only in Cluster 1 (Fig. S4A,C,D). Cluster 5 is enriched for protein folding genes, Cluster 4 for stress response and catabolic processes and Cluster 3 for genes associated with mitochondrial aerobic respiration (Table S1).

**Figure 4.**
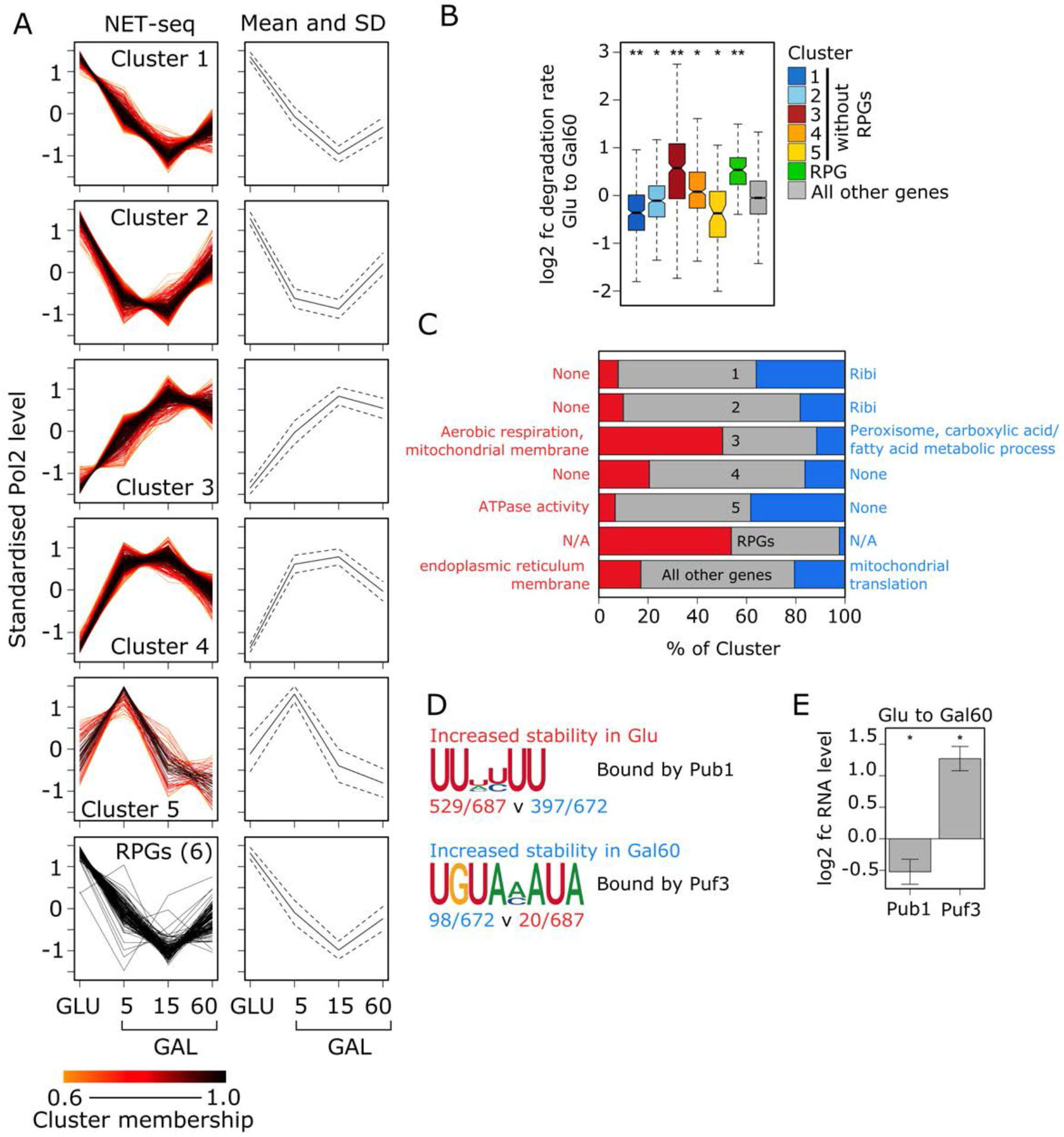
Altered transcript degradation rates fine tune transcript levels in response to stress. A) Standardized profiles of mean regularized log2 transformed Pol2 levels at genes determined from NET-seq samples from cells grown exponentially in YPD (Glu) and after 5 (Gal5), 15 (Gal15), and 60 min (Gal60) following the replacement of YPD for YPG. Genes were clustered into 5 clusters according to the shape of their profiles. Only genes showing a significant 2-fold change in Pol2 levels between a timepoint and any subsequent timepoint in this time series were included in the clustering. Only genes with a membership score greater than 0.6 for a particular cluster are shown. Profiles are coloured according to their cluster membership score. B) Boxplots showing the log2 fold change in degradation rate from Glu to Gal60 for each of the Pol2 profile-determined clusters with the exception of RPGs which have been placed in a separate cluster. Asterisks mark mean cluster log2 fold changes significantly (*, p < 0.05; **, p < 1×10^−10^, Welch’s t-test adjusted for multiple testing (Holm method)) different from that of genes not in any cluster. C) Proportions of genes in each cluster showing a decrease, no change or increase in mRNA stability from Glu to Gal60 (according to the thresholds detailed in A). Only genes showing a significant change in RNA level between any of the Glu, Gal5, Gal15 and Gal60 conditions are included. Gene ontology analysis was carried out on each of the proportions showing a decrease or increase and key terms are displayed to the side of these proportions. D) Sequence logos of motifs identified by DREME (Bailey et al., 2011) as enriched in the 3’UTR of mRNAs showing increased stability in Glu (top) or increased stability in Gal60 (bottom). mRNAs showing the opposing change in stability were used as control sequences. Number of mRNAs with the motif in test and control sequences are shown. Only 3’UTRs longer than 50 nt were included. RNA binding proteins (RBPs) which potentially bind the motifs are shown. E) Bar chart showing the log2 fold change in RNA level from Glu to Gal60 for the two RBPs detailed in E. Asterisks mark significant changes (p < 0.05, Wald test, adjusted for multiple testing using the Benjamini–Hochberg method). Error bars show the standard error. See also Figure S4 and Table S1.

### Genes with altered transcript stability have Pub1 or Puf3 binding sites in their 3’ UTR

As transcript degradation rates correlate with Pol2 levels and synthesis rates (23,25), we analysed how degradation rates change for each of our NET-seq data-determined clusters (Fig. 4B), exclusive of RPGs, which were included as an extra cluster consisting of 131 RPGs identified by Jorgensen et al. (2004) (45). For Clusters 1 to 4, where there is either an increase or decrease in transcription there is also a significant increase or decrease, respectively, in the average degradation rate (Fig. 4B). This change is on average greater for those clusters which are slower to reach there minimal (Cluster 1 v 2) or maximal (Cluster 3 v 4) Pol2 levels. Interestingly, despite showing a similar transcription profile to Cluster 1 genes, RPGs show an increased average degradation rate.

We applied a cut off to determine genes showing a log2 fold change in degradation rate greater than +/-0.5 as well as a significant change in RNA level at least at one timepoint during the 1h time series upon the switch to Gal. We intersected these genes with the NET-seq clusters and carried out GO analysis to determine enriched terms in the different groups (Fig. 4C). Mitochondrial genes for aerobic respiration in cluster 3 are enriched in the group of genes showing increased degradation while mitochondrial RPGs, which are not enriched in any of the NET-seq clusters, show an enrichment in the group of genes showing decreased degradation.

The 3’UTR of mRNAs has been shown to contain motifs which are bound by a range of RNA binding proteins which can modulate RNA stability. We carried out DREME motif enrichment analysis to identify motifs in either group of genes showing an increase or decrease in transcript stability in Gal60. For gene transcripts showing an increase in stability in Gal60, there was an enrichment for the motif with consensus UGUAMAUA (Fig. 4D), which has been shown to be bound by Puf3 (46). Puf3 has been shown to promote RNA degradation during growth in glucose (47). In response to a switch to glucose-depleted media, Puf3 is phosphorylated and targets its bound transcripts to ribosomes for translation instead of degradation (48). Here we also show that Puf3 shows significantly increased RNA levels following the switch to galactose which may increase its ability to prevent RNA degradation of this cohort of transcripts (Fig. S4E). For genes showing a decrease in stability in Gal60 (increased stability in Glu), there was an enrichment for a U-rich motif (Fig. 4D). These are potentially bound by Pub1, which plays a role in protecting transcripts from degradation (49,50). The decrease in Pub1 transcript levels seen after 60 min in YPG (Fig. S4E), may lead to a decrease in this protective effect and increased degradation of these transcripts.

### Genes with different kinetics of induction and repression are associated with specific transcription factors

Transcription factors are proposed to play a key role in the response to stress. Key factors, such as Msn2 and Sfp1, show differential association with specific promoters as a result of stress, often resulting from altered phosphorylation and subcellular localisation (7,17,33,45,51-57). Msn2 associates with the consensus motif (RGGGG) in the promoters of many genes showing an increase in transcript levels on stress (58). By contrast, RiBis, which are repressed on stress, have rRNA processing element (RRPE; TGAAAAWTTTY) and polymerase A and C (PAC; GCGATGAGMT) motifs in their proximal promoter, bound by the transcription factors Sfp1 and Dot6/Tod6, respectively. Sfp1 is a transcriptional activator that is nuclear located during exponential growth and is phosphorylated and translocated into the cytoplasm in response to a broad range of stresses (45). In contrast, Dot6 and its paralog Tod6 are repressors that are activated in response to nutrient deprivation (59). RPGs are activated by the binding of Rap1 to the distal promoter region in optimal conditions (60).

We examined for significant enrichment of binding motifs for Rap1, Dot6/Tod6, Sfp1, Msn2 or Hap5 in the distal or proximal promoters of genes in clusters 1-5 and the RPGs (Fig. 5A,B). With the exception of Cluster 5, which despite showing rapid induction is neither enriched nor depleted for any motifs, the remaining clusters contained motifs for Msn2, Sfp1 and Dot6 which corresponded with whether they were repressed or induced in Gal (Fig. 5C). Given the distinct regulatory programme involved in RPG expression (9), we excluded RPGs from the five NET-seq clusters and confirmed enrichment with Rap1 consensus motif in the distal promoters (Fig. 5A).

**Figure 5.**
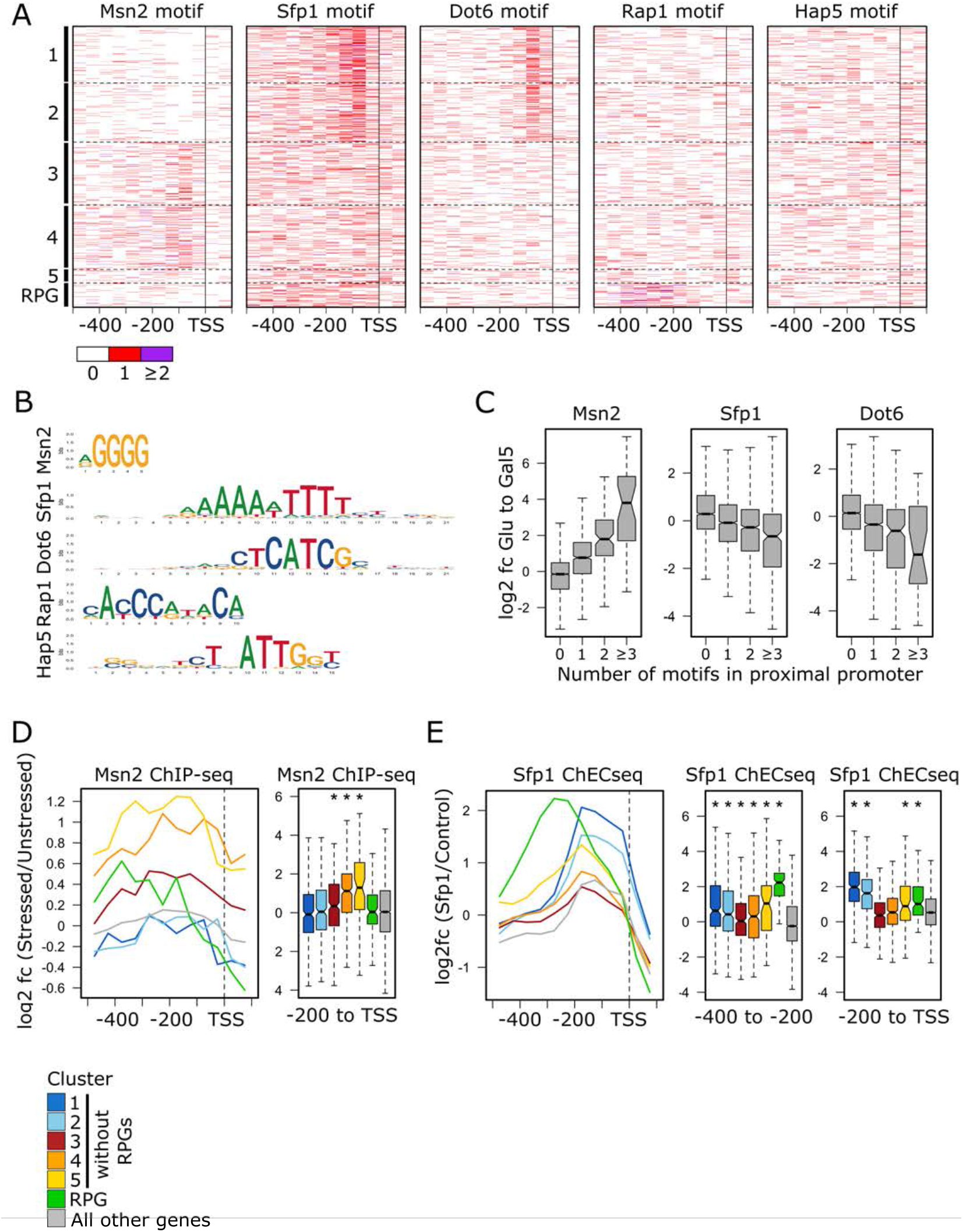
Genes with different kinetics of induction and repression are associated with specific transcription factors. A) Heatmaps showing the number of motifs in each 50 bp window within the promoter region of the genes in each Pol2 profile-determined cluster. The 131 RPGs (45) are excluded from each cluster and form their own cluster. Motifs can occur on either strand apart from the Sfp1 motif, which displays directionality. 50 bp window assignment is determined from the position of the central nucleotide of the motif. B) Sequence logos of the motifs. C) Boxplots showing the log2 fold change in Pol2 level between Glu and Gal5 conditions at genes with different numbers of motifs in their promoter proximal region (TSS-200 nt to TSS). D) Mean log2 fold change in Msn2 ChIP-seq signal in *S. cerevisiae* cells after 20 min exposure to glucose-starvation stress (34) in 50 bp windows across the promoter region of genes for each cluster (left panel). Boxplots showing the log2 fold change in this signal in the promoter proximal region (TSS-200 nt to TSS) for genes of each cluster (right panel). Asterisks mark mean cluster log2 fold changes significantly (p < 0.05, Welch’s t-test adjusted for multiple testing (Holm method)) greater than that of genes not in any cluster. E) Mean log2 fold change in Sfp1 ChEC-seq signal relative to control ChEC-seq signal in *S. cerevisiae* cells growing exponentially in glucose-rich media (33) in 50 bp windows across the promoter region of genes for each cluster (left panel). Boxplots showing the log2 fold change in this signal in the promoter distal (TSS-400 nt to TSS-200 nt, middle panel) and proximal (TSS-200 nt to TSS, right panel) region for genes of each cluster. Asterisks mark mean cluster log2 fold changes significantly (p < 0.05, Welch’s t-test adjusted for multiple testing (Holm method)) greater than that of genes not in any cluster. Boxplots (C-E) show the median (central line), median +/-interquartile range (IQR)/square root (number of genes) (notch), IQR, (box edges), most extreme data point no more than 1.5× IQR from box edges (whiskers). Outliers outside the whiskers are not shown. See also Figure S5.

As this analysis is based solely on motif analysis, we asked whether the motifs in these clusters are protein bound or not by analysing published data for chromatin associated Msn2 (Fig. 5D, Fig. S5A) (34) or Sfp1 (Fig. 5E, S5A) (33). After 20 min exposure to glucose starvation conditions, Msn2 shows significantly increased average binding to the promoters of Cluster 3 and 4 genes with greater binding to Cluster 4 than 3 (Fig. 5D). In addition, overexpression of Msn2 leads to increased RNA levels of both Cluster 4 and 3 with Cluster 4 RNA levels rising on average earlier than those of Cluster 3 (Fig. S5B). Conversely, exposure of a strain lacking *msn2* and *msn4* (a partially redundant paralog of *msn2*) to H_2_O_2_ stress results in reduced induction of Cluster 3 and 4 transcripts without the earlier induction of Cluster 4 over 3 transcripts seen in a wildtype strain (35) (Fig. S5B).

Interestingly, despite Cluster 5 not showing enrichment for Msn2 binding motifs (Fig. 5A), Msn2 also shows increased binding to Cluster 5 gene promoters upon stress (Fig. 5D). This suggests that Msn2 may bind to these promoters indirectly via interaction with another factor. In support of this, overexpression of Msn2 does not lead to an increase in Cluster 5 RNA levels (Fig. S5B). However, loss of both Msn2 and Msn4 does not affect Cluster 5 gene induction in response to H_2_O_2_ stress (35), suggesting that Msn2/4 may be redundant with other factors or not involved in the induction of these genes (Fig S5B). Alternatively, its role may be specific to particular types of stress. This highlights the complex nature of transcription factor binding to promoters.

Chromatin endogenous cleavage sequencing (ChEC-seq) (33) reveals significant enrichment of the Sfp1 signal in Clusters 1, 2 and 5 in the promoter proximal region, and, possibly via its indirect recruitment by Rap1, high significant enrichment in the distal promoter region of RPGs (7,51,61,62) (Fig. 5E). Cluster 1 shows the higher average number of Sfp1 motifs and higher ChEC-seq signal than Cluster 2 in this region. This may explain why these genes are more highly expressed on average during exponential growth compared to Cluster 2 genes and take longer to reach their minimal expression level during the switch to galactose. Although Alberts et al. (2019) (33) did not report Sfp1 binding to RPGs by ChEC-seq, our reanalysis of their data does show significant enrichment of Sfp1, particularly over the distal promoter region (Fig. 5E). Sfp1 overexpression leads to increased Pol2 levels in Cluster 1, 2 and RPG genes but no effect on Cluster 5 genes despite the significant enrichment of Sfp1 ChEC-seq signal in the promoters of these genes (Fig. S5A). Removing Sfp1 by anchor away leads to a decreased expression at Clusters 1 and 2 and at RPGs (Fig. S5C).

We next applied motif enrichment analysis using DREME (36) to identify additional enriched motifs in particular clusters. This revealed enrichment for an RCCAATCA motif in Cluster 3 which is bound by Hap2/3/4/5 transcription factors known to be involved in the induction of genes for aerobic respiration (Fig. 5A,S5A). Cluster 5 shows enrichment for motifs bound by the transcriptional activator Hsf1 and repressor Rgt1. It is interesting to note that both Sfp1 and Msn2 exhibit binding to Cluster 5 promoters (Fig. 5D, E) despite lacking enrichment for their binding motifs (Fig. S5A). This may suggest that Cluster 5 promoters are more accessible to transcription factors in general, which may assist the rapid induction and subsequent repression kinetics observed for these genes. This prompted us to examine how transcription elongation and nucleosome positions change during the Glu to Gal shift.

### Transcription elongation is disrupted with a global increase in the level of Pol2 in the 5’ gene region relative to that in the gene body

In addition to revealing the total level of Pol2 over every gene during stress, NET-seq also reveals the profile of transcriptionally engaged Pol2 along genes at single nucleotide resolution, capturing the elongating, stalled and backtracked enzyme, with Pol2 levels generally higher in the promoter proximal region than in the gene body (18,22,42). We analysed how the Pol2 profiles change following the switch to Gal, showing that Pol2 shows even more increased levels in the 5’ region of the gene relative to the gene body within 5 min of the switch (Fig. 6A). This shift is particularly evident when we plot the average profiles of genes, after excluding very highly or poorly expressed genes (Fig. 6B; S6A,B). This is reflected in the travelling ratio (TR) of Pol2 after the switch to galactose (Fig. 6C; S6C,D).

**Figure 6.**
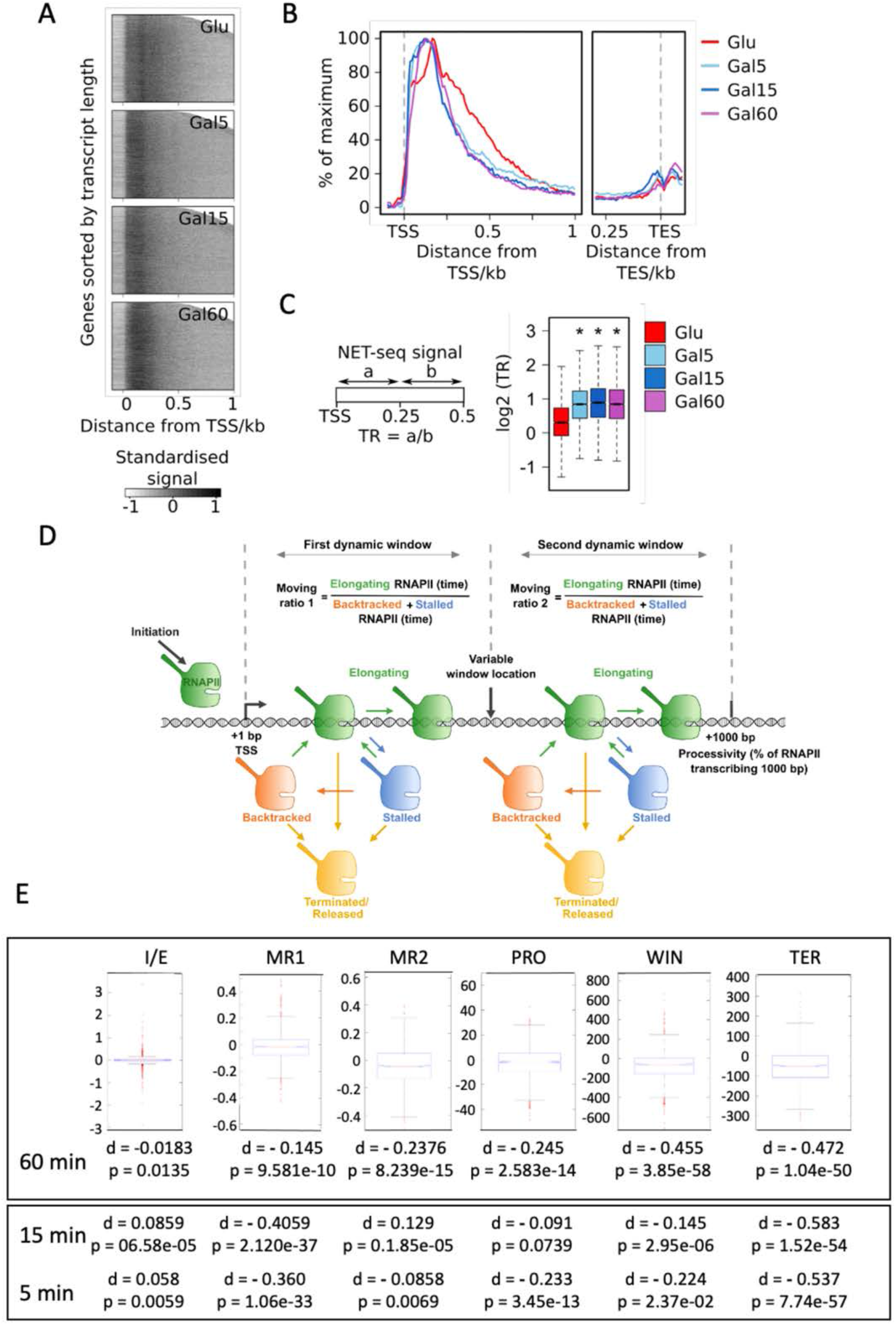
Transcription elongation is disrupted with an increase in the level of Pol2 in the 5’ gene region relative to that in the gene body, irrespective of any change in total Pol2 levels. A) Heatmaps showing the Pol2 level profile across all genes from TSS-100 nt to TSS+1000 nt for Glu, Gal5, Gal15 and Gal60 conditions. NET-seq data from combined biological repeats has been averaged in 10 nt bins across genes. Bins with signal less than 0 are set to 0. Bins are then standardized for each gene by subtracting the mean and then dividing by the standard deviation of bins within the window TSS to TSS+1000 nt. B) Metagene Pol2 level profile showing the average standardized Pol2 profile for each condition for all genes. For profiles around the TSS, bins within the last 200 nt of genes are excluded. For profiles around the TES, bins within the first 500 nt of genes are excluded and genes shorter than 600 nt are excluded. The top and bottom 5 % of bins are excluded prior to averaging. Each profile is plotted as a percentage of the maximum signal. C) Schematic detailing the calculation of the travelling ratio. For each gene, the mean of the bins in the window from TSS to TSS+250nt is divided by the mean of the bins in the window from TSS+250nt to TSS+500nt. Boxplots showing the travelling ratios for each condition for all genes. D) Schematic of the mathematical model. Model describes RNAPII transcription reaching 1000 bp with initiation rate (initiation per min), elongation rate (kb per min), stalling, backtracking, termination (TER) events (determined by Poisson distribution), variable window (WIN) location (bp). Moving ratio 1 and 2 (MR) describes number of RNAPII elongating compared to backtracked or stalled RNAPII within respective transcription window. Processivity (PRO) indicates % of RNAPII reaching 1000 bp. E) Boxplots of the pairwise differences in inferred parameters for each gene for Gal60. Metrics were obtained for each gene in Glu, Gal5, Gal15 and Gal60 and each Gal condition quantitatively compared to Glu. The significance of the changes was reported by calculating p-values and the magnitude of the changes was reported by calculating Cohen’s d (88). Cohen’s d is computed by taking the mean difference between the Glu and Gal metric value divided by the standard deviation of the differences. The value of Cohen’s d gives a measure of the effect size of the change (88). Positive and negative values indicate a relative increase or decrease in the given metric, respectively. See also Figure S6.

We used our model (63) (Fig. 6D) to computationally simulate Pol2 dynamics by considering initiation, elongation, occlusion of Pol2 by a downstream Pol2, collision of Pol2s, stalling, backtracking, resolution of collision/backtracking/stalling events, and early termination. In contrast to previous approaches (64-66), we set two distinct windows of transcription in which stalling and backtracking parameters can be different, motivated by the notion that Pol2 is subject to distinct regulation in the early and late stages of transcription (67). The Pol2 position provided by experimental NET-seq data for cells in glucose or after 5, 15 and 60 min in galactose at single nucleotide resolution was fitted to the shape of the transcription profile simulated by the model (Fig. S6E,F). Modelling derives six key metrics that can be inferred from the shape of the WT RNAPII distribution: 1) ratio of the rate of initiation compared to elongation (I/E), 2) ratio of RNAPII moving compared to stalled or backtracked (moving ratio) in window 1 (MR1), 3) the size of window 1 (WIN), 4) the mean location of early termination (TER), 5) the moving ratio in window 2 (MR2), and 6) the processivity of RNAPII (% of initiating polymerase reaching 1000 nt; PRO) (Fig. 6D). To test the extent of the change in each metric, the parameter values were obtained for each gene in Glu and Gal60 and the two conditions were quantitatively compared. The difference values (Cohen’s **d** which gives an effect size for the comparison between two means) between the 6 metrics revealed by the model indicate those most affected after 60 min in Gal compared to Glu and, using the same model, at 5 and 15 min (Fig. 6E). The metric that changes consistently at all three time points is the early termination location moving closer to the promoter (d = - 0.53, -0.58, -0.47 at 5, 15 and 60 min, respectively). The differences in other metrics such as window location, processivity, and moving ratio 1, change over time suggesting that elongation by Pol2 is constantly adapting to the stress, although we note that the initiation to elongation ratio does show a significant difference at any time point. This analysis suggests that stress affects transcription elongation dynamically, broadly and globally, leading to earlier termination of transcription regardless of whether transcription is induced or repressed. We note, however, that the NET-seq profiles at induced or repressed genes are subtly different, particularly in the promoter proximal region and may reflect distinct forms of elongating Pol2 (Fig. 7A,B).

**Figure 7.**
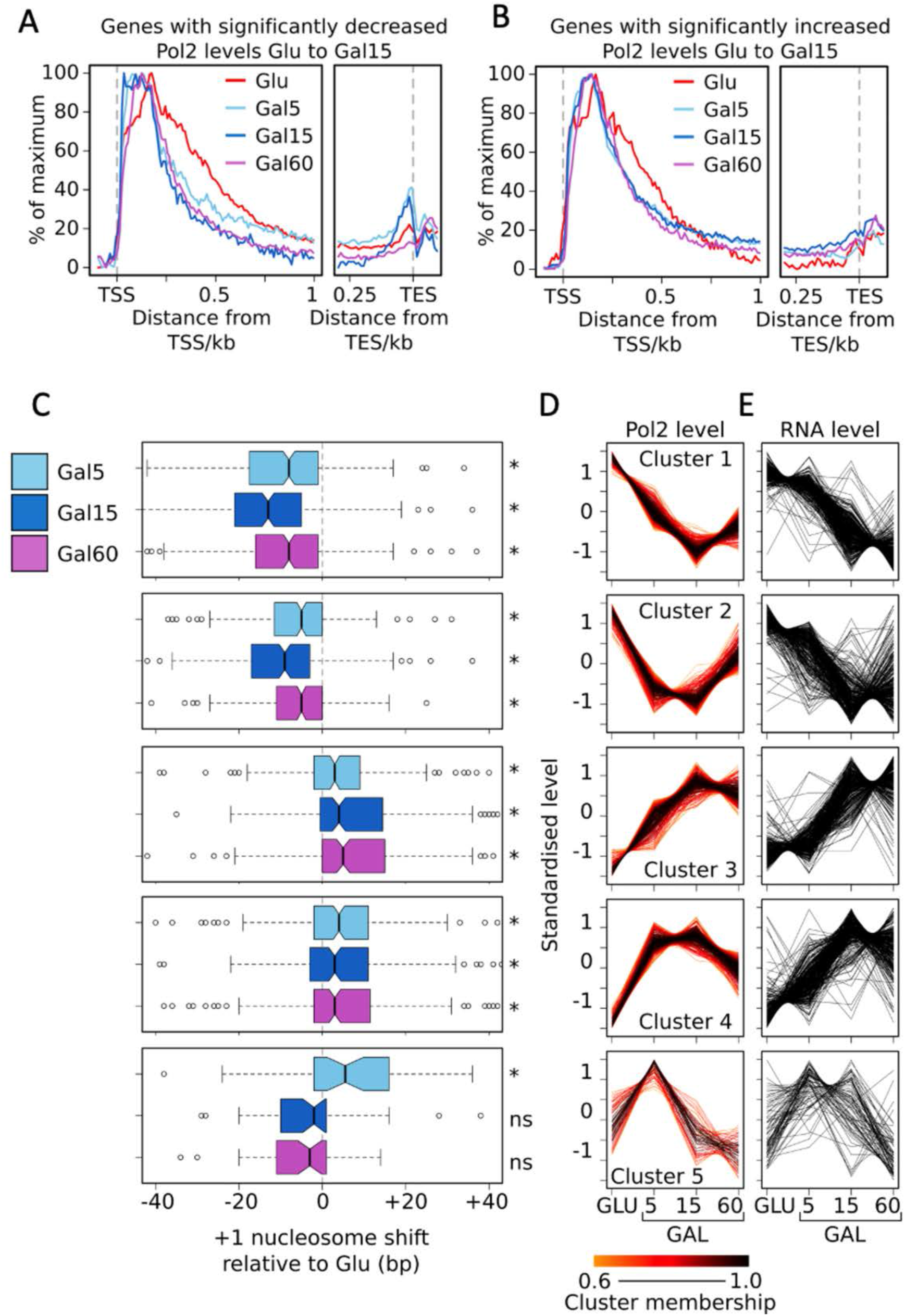
Plus 1 nucleosomes shift downstream or upstream as Pol2 levels increase or decrease, respectively. A and B) Metagene Pol2 level profiles showing the average standardized Pol2 profile for each condition for all genes showing a significant decrease (A) or increase (B) in Pol2 level from Glu to Gal15. For profiles around the TSS, bins within the last 200 nt of genes are excluded. For profiles around the TES, bins within the first 500 nt of genes are excluded and genes shorter than 600 nt are excluded. The top and bottom 5 % of bins are excluded prior to averaging. Each profile is plotted as a percentage of the maximum signal. C) Standardized profiles of mean regularized log2 transformed Pol2 levels at genes determined from NET-seq samples from cells grown exponentially in YPD (Glu) and after 5 (Gal5), 15 (Gal15), and 60 min (Gal60) following the replacement of YPD for YPG. Genes were clustered into 5 clusters according to the shape of their profiles. Only genes showing a significant 2-fold change in Pol2 levels between a timepoint and any subsequent timepoint in this time series were included in the clustering. Only genes with a membership score greater than 0.6 for a particular cluster are shown. Profiles are coloured according to their cluster membership score. D) Standardized profiles of mean regularized log2 transformed RNA levels determined from RNA-seq samples from Glu, Gal5, Gal15 and Gal60 conditions for genes of the Pol2 profile-determined clusters. E) Boxplots showing the shift in bp in the position of the +1 nucleosome dyad in Gal5, Gal15 and Gal60 conditions relative to the position in Glu for genes of the Pol2 profile-determined clusters at genes clustered according to their patterns of induction and repression. Asterisk mark significant (p < 0.05, Mann-Whitney test) average shifts.

### The position of the +1 nucleosome shifts either downstream or upstream as Pol2 levels increase or decrease, respectively

Having established that widespread transcriptional changes occur within 5 min of a switch to Gal, and the role of transcription factors in the rapid induction or repression of transcription at different gene clusters, we aimed to assess how chromatin structure changes in relation to these transcriptional changes. To do this we profiled nucleosome positions by MNase-Seq at Glu, Gal5, Gal15 and Gal60 time points. We then determined how the position of the +1 nucleosome changes for genes from our five NET-seq clusters (Fig. 7E) and compared this with Pol2 (Fig. 7C) and RNA (Fig. 7D) levels at each timepoint in each cluster. Within 5 minutes, for all clusters showing a decrease in Pol2 levels (clusters 1 and 2) there is a corresponding upstream shift in the position of this nucleosome compared to Glu. Conversely, for all clusters showing an increase in Pol2 levels (clusters 3-5) within this short time period, there is a corresponding downstream shift in the position of this nucleosome. For cluster 5, which shows a return in Pol2 levels to that seen in Glu after 15 min, the +1 nucleosome also returns to its position in Glu at this time point. As the +1 nucleosome abuts the TSS, an upstream nucleosome shift may repress the ability of Pol2 to initiate transcription from this site while a downstream shift may enable it. As the changes in Pol2 levels and nucleosomes positions are so dynamic, without clustering and the direct analysis of Pol2 levels giving us high spatial and temporal resolution, we would not have been able to uncover these correlations. This is particularly evident for transcript levels, which are generally temporally disconnected from Pol2 levels and changes in +1 nucleosome changes (Fig. 7C,D,E). Taken together these data support coordinate gene-specific changes to transcription factor occupancy, nucleosome positioning and Pol2 occupancy that reflect dynamic activation or repression of transcription and subsequent changes in transcript levels in response to this carbon source stress. In addition, the global response of Pol2 to stress, leading to early termination of transcription in the promoter-proximal window, is distinct from the gene-specific changes in +1 nucleosome positioning.

## DISCUSSION

By using a system with high temporal and spatial resolution and controlling for the stress imposed on the yeast during the experimental procedures, this work uncovers new and mechanistically distinct features of the acute nuclear stress response in yeast which has two distinct components. We uncover a genome-wide effect on Pol2 occupancy that leads to an upstream shift in the peak of accumulation that is normally focused about +170 nucleotides from the TSS just upstream of the +2 nucleosome. Modelling supports this shift resulting from early termination of transcription and is unrelated to whether Pol2 occupancy or transcript levels increase or decrease, and so is also independent of +1 nucleosome dynamics. The second, distinct component to the acute nuclear stress response is the highly dynamic patterns of induction and repression of transcription at specific subgroups of genes related to TF occupancy and the dynamic repositioning of the +1 nucleosome. This supports two distinct effects of glucose depletion on yeast transcription, one general focused on Pol2 and one gene-specific focused on transcription factors and the +1 nucleosome.

Unravelling these relationships relied on base-pair resolution data and the ability to sample precisely at early time points using NET-Seq. The lack of strong correction between nucleosome dynamics, transcript levels and nascent transcripts in experiments using metabolic labelling is likely to reflect the lack of temporal resolution. For example, it takes 6 minutes for thiouracil (4tU) to be taken up and metabolised into a form where is can be incorporated into newly synthesised transcripts (25), although it is an excellent technique for other applications. χCRAC (68) of Pol2 has similar temporal resolution to NET-seq but due to the necessity to cross-link Pol2 to transcripts, lacks the dynamic range of signal afforded by NET-seq. Moreover, the depth and single nucleotide resolution of the data sets enables the shape of NET-seq profiles to be analysed by modelling and simulation (24,63), defining which of the six key metrics that can be inferred from the shape of the Pol2 distribution in unstressed cells are changed on acute stress. The changes involve a shift in the peak of Pol2 accumulation at 170 nt towards the promoter, and changes in the signal over the gene body and at the polyadenylation site. Interestingly, the genome-wide component of the response occurs regardless of whether Pol2 occupancy over genes increases, decreases or remains unchanged (69) (this study). The effects on Pol2 may be direct or mediated by elongation factors such as FACT (70), Spt4/5 (63) or Paf1 (26) or other, as yet less well understood, components of the regulated transcriptional response and the feedback control from the cytoplasm to buffer transcription and transcript levels (23,25,71-75).

Our data show a very clear relationship between transcription and nucleosome dynamics both for genes whose expression increases and those whose expression decreases. Due to many techniques lacking temporal resolution or relating nucleosomes to transcript levels, these relationships have not been observed at a genome-wide scale before although are widely predicted to be a key component of gene regulation (9). These shifts are relatively small, and highly dynamic, showing maximal movement with the point at which transcription, assessed by Pol2 occupancy, is maximally induced or repressed. Whether these changes are controlled by active chromatin remodelling by ATPases such as Isw1, Isw2, RSC or Swi/Snf (76-78), recruited via stress-related transcription factors or reflect the remodelling activity of Pol2 itself is not clear at present. This dynamic relationship between the onset of transcription and nucleosome positions would not have been evident had nucleosome positions been related to changes in transcript levels, as these are temporally offset early in the acute stress response (68) (this study), or if the system used lacked the necessary temporal resolution.

The relationship between gene expression and acute stresses such as heat shock, H_2_O_2_ stress, salt shock or glucose starvation have been assessed in yeast using cross-linking techniques such as ChIP-seq (34,79), ChIP-exo (69), χCRAC (68) of Pol2, or metabolic labelling techniques such as GRO-seq (80) or comparative dynamic transcriptome analysis (cDTA) (25,79). ChIP of Pol2 lacks spatial resolution and fails to find strong correlations (>0.7) between either transcript levels or metabolically labelled nascent transcripts, 12 and 24 min after imposition of stress (79), although recent ChIP-exo data suggests the accumulation of transcriptionally engaged Pol2 on H_2_O_2_ stress, known as a stall, is focused at the +2 nucleosome (69). Spt4 in the Spt4/5 complex is required for transcription through the +2 nucleosome (63). We expected the effect of the glucose starvation used here mimic the effects of loss of Spt4 or H_2_O_2_ stress on polymerase during early elongation. Although we did see a global response, this was mechanistically focused on earlier transcription termination and did not reveal a transcription elongation defect in the promoter proximal region as we observed in strains lacking Spt4 (63). This suggests that there is not one global response to stress but rather each stress is sensed and signals to different aspects of the machinery that control the global transcriptional response.

With respect to the gene-specific response to an acute stress, our data are in line with those reported by others (69). A key question is how the genome-wide response evident in the shape of the metagene profiles is distinct from the gene-specific increases or decreases in Pol2 occupancy. One possibility is that transcription factors determine the acute response in terms activation or repression of transcription while the global changes affect the actual process of transcription by Pol2. As we are only assessing transcription and transcript levels, it is not clear how the global changes will impact on downstream events such as subcellular localisation of transcripts (26,72,81) and their translation, which also responds to stress (82,83) and how much this impacts overall protein levels, particularly in the early stages of the acute stress response when transcript and translation levels change dramatically but do not always reflect acute changes in protein levels (84-87).

## DATA AVAILABILITY

### ACCESSION NUMBERS

NET-seq data for Glu, Gal5, Gal15 and Gal60 were obtained from ArrayExpress E-MTAB-4568

All sequencing data generated in this study are deposited at GEO (https://www.ncbi.nlm.nih.gov/geo/) under accession code GSE154184. The reviewer token is **arozoquixdwtncz**

## SUPPLEMENTARY DATA

Supplementary Data are available at NAR online.

## ACKNOWLEDGEMENTS

H.F. and J.M. designed the experiments. H.F. performed experiments and analysed datasets. T.B. and A.A. performed and analysed mathematical modelling. J.M. and H.F. wrote the manuscript, with input from all authors.

