## Supplementary material for "Global and Gene-specific Transcriptional Responses to Acute Stress": Supplem

**Figure S1 Galactose metabolising genes do not show large increases in expression 60 min after the transfer of cells from glucose-to-galactose-containing media**

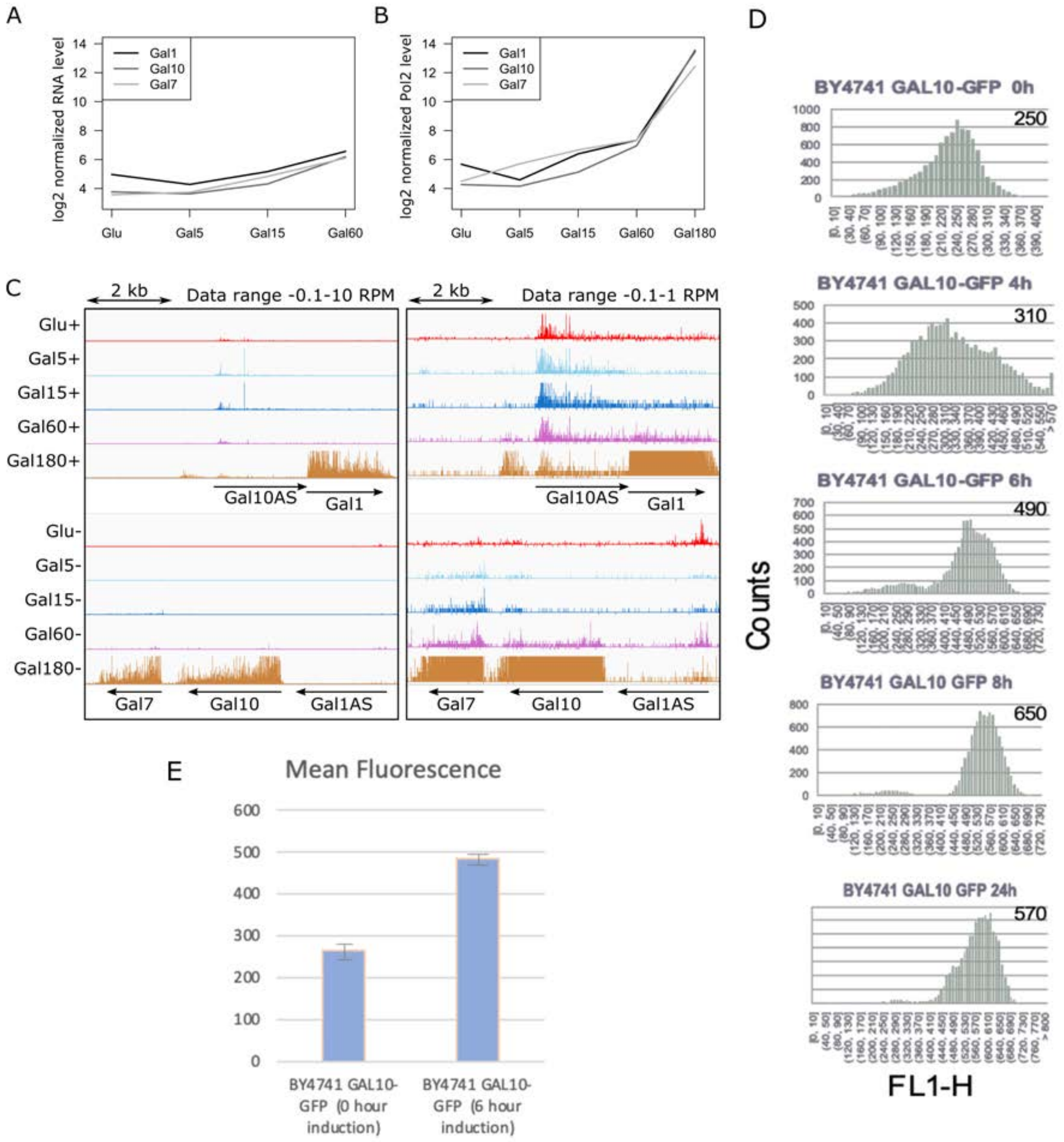

A and B) Mean regularized log2 transformed RNA (A) and Pol2 (B) levels at *GAL1*, *GAL7* and *GAL10* genes as determined from RNA-seq and NET-seq samples, respectively, from cells grown exponentially in YPD (Glu) and after 5 (Gal5), 15 (Gal15), 60 min (Gal60) and 180 min (Gal180, Pol2 levels only) following the replacement of YPD for YPG. C) NET-seq data for both strands showing counts per million counts (CPM) for Glu, Gal5, Gal15, Gal60 and Gal180 samples at the locus containing *GAL1*, *GAL7* and *GAL10* genes. All tracks show the combined counts from 2 biological replicates except Gal180 (1 biological replicate). Data range -0.1-10 CPM (left panel) -0.1-1 CPM (right panel). Panels were prepared using Integrative Genomics Viewer (IGV). D) FACS analysis of GAL10-GFP after 0,4,6,8 and 24 hours in galactose showing count distributions, with maximum peak intensities marked for each time point. E) Mean fluorescence (GAL10-GFP) before and after 6 h induction in galactose. Error bars show the standard error.

**Figure S4 Further characterisation of the gene clusters**

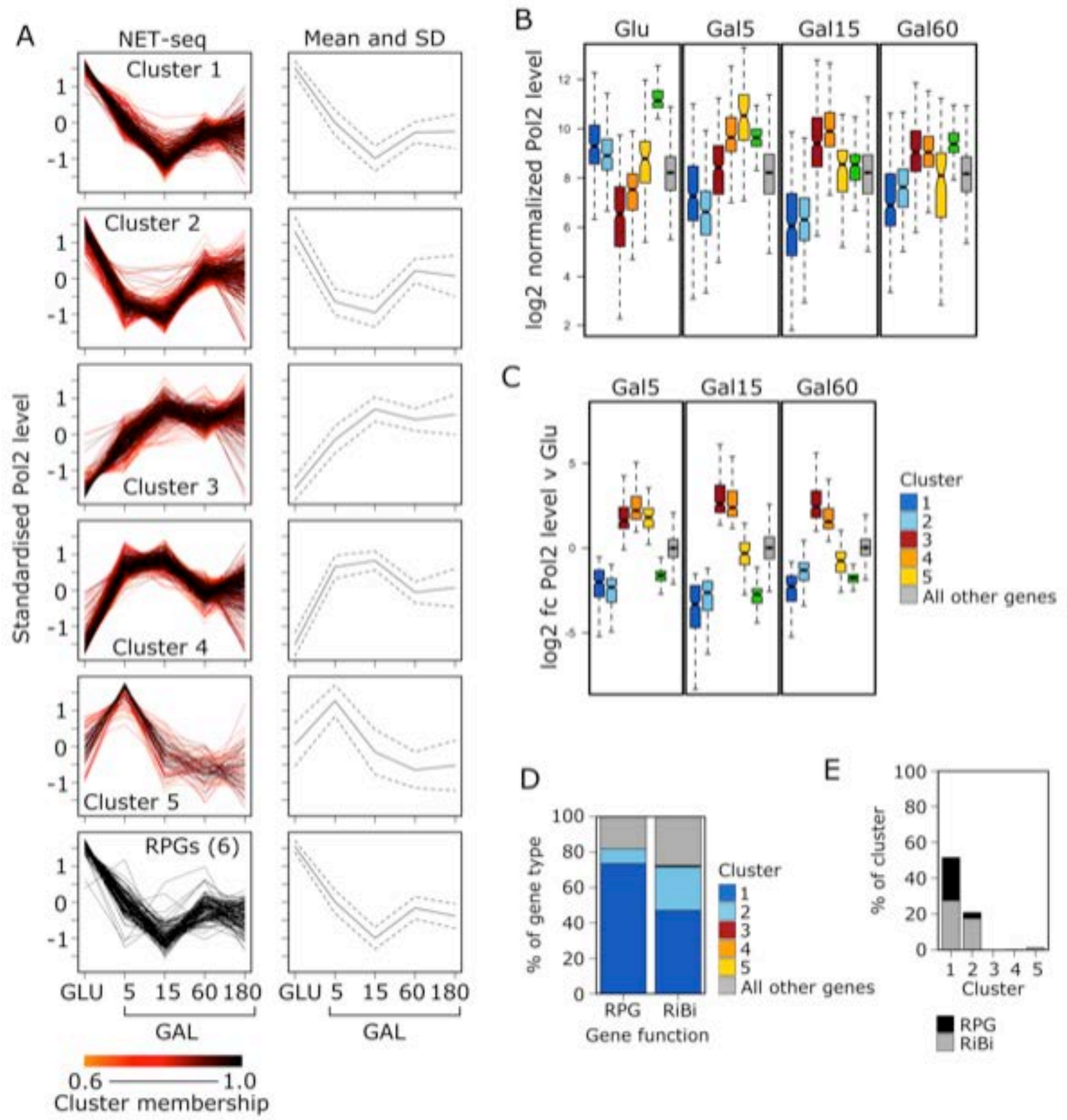


A) Standardized profiles of mean regularized log2 transformed Pol2 levels at genes determined from NET-seq samples from cells grown exponentially in YPD (Glu) and after 5 (Gal5), 15 (Gal15), 60 (Gal60) and 180 min (Gal180) following the replacement of YPD for YPG. Left panels show individual gene profiles coloured according to their membership score for that cluster (RPG traces are not coloured). Right panels show the mean (solid line) and +/-1 standard deviation away from the mean (dashed line) of the individual gene profiles of each cluster. B) Boxplots showing the log2 fold change in Pol2 levels for genes in each cluster for the conditions shown relative to those in Glu. D) Percentage of gene type of each cluster that are RPGs or RiBis. E) Percentage of each cluster that are either RPGs or RiBis.

**Figure S5 Genes with different kinetics of induction and repression are associated with specific transcription factors**

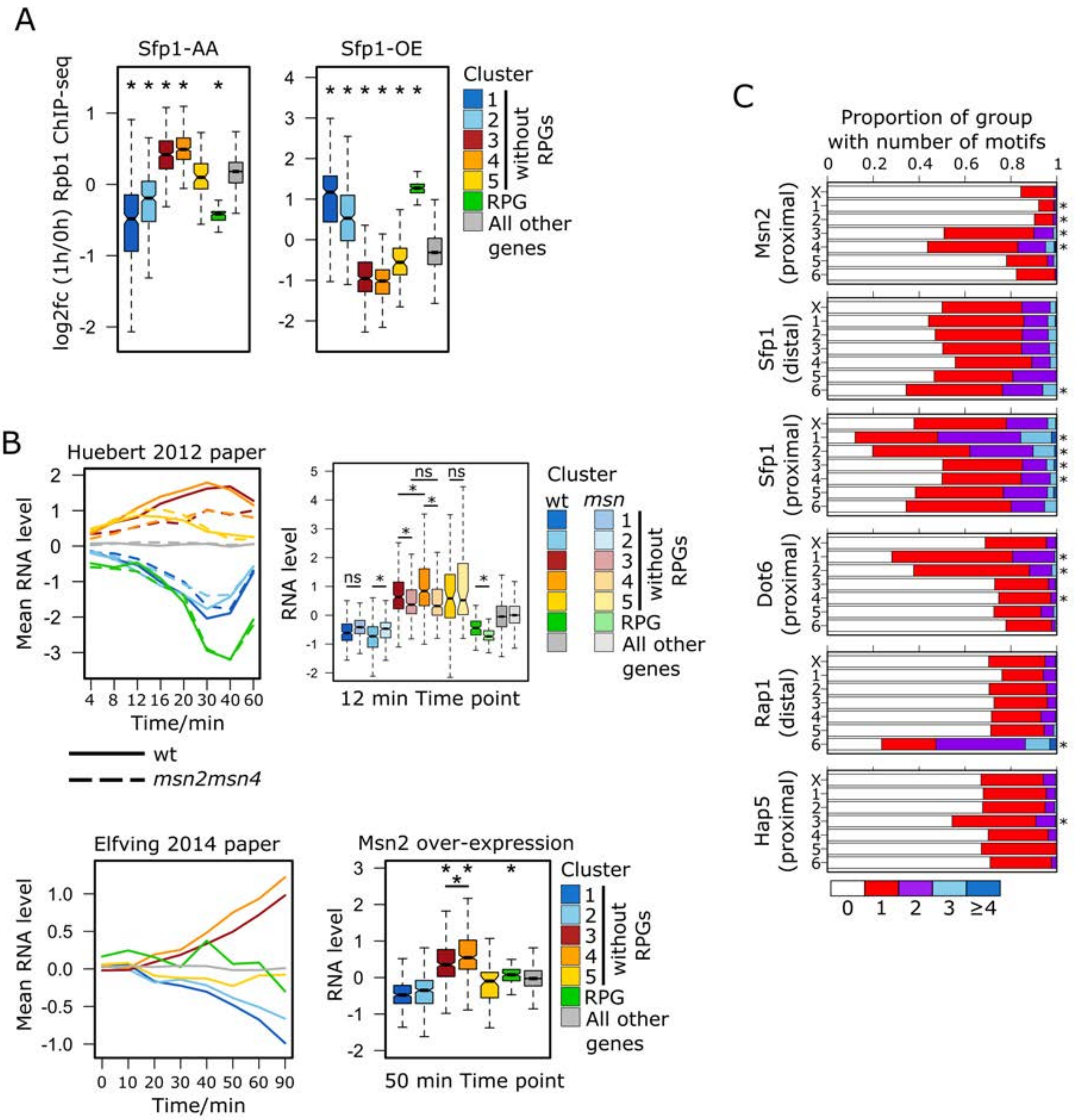


A) Boxplots showing the log2 fold change in Rpb1 ChIP-seq signal in *S. cerevisiae* cells after either Sfp1 anchor away (left panel) or Sfp1 overexpression (right panel) for 1 h {Albert, 2019 #18990} for each cluster. Asterisks mark mean cluster log2 fold changes significantly ($p < 0.05$, Welch's t-test adjusted for multiple testing (Holm method)) different from that of genes not in any cluster. B) Mean RNA level at time points following exposure of wild type (solid lines) and *msn2msn4* double knockout mutant (dashed lines) *S. cerevisiae* cells to hydrogen peroxide stress (29) for each cluster (left panel). Boxplots showing the RNA level at the 12 min time point for genes of each cluster for both wildtype and mutant strains. Welch's t-tests were used to test the significance (*, $p < 0.05$; ns, $p >= 0.05$) of any difference between the mean wildtype and mutant RNA level for each cluster. P values were adjusted for multiple testing (Holm method). Welch's t-tests were also used to test the significance (*, $p < 0.05$; ns, $p >= 0.05$) of any difference between the mean RNA level of clusters 3 and 4 in either the wildtype or mutant strains. Mean RNA level at time points following Msn2 over expression in *S. cerevisiae* cells (28) for each cluster (left panel). Boxplots showing the RNA level at the 50 min time point for genes of each cluster. Asterisks mark mean cluster RNA levels significantly ($p < 0.05$, Welch's t-test adjusted for multiple testing (Holm method)) greater than that of genes not in any cluster. A Welch's t-test was also used to test the significance of any difference between the mean RNA level of clusters 3 and 4. C) The proportion of genes with particular numbers of each motif in either the promoter distal (TSS-400 nt to TSS-200 nt, middle panel) or proximal (TSS-200 nt to TSS, right panel) region for each Pol2 profile-determined cluster. The 131 RPGs (35) are excluded from each cluster and form their own cluster (cluster 6). Asterisks mark clusters with significantly ($p < 0.05$, two sample Poisson Rate test adjusted for multiple testing (Holm method)) different numbers of motifs relative to cluster X (all other genes).

**Figure S6 Transcription elongation disruption is observed across replicates**

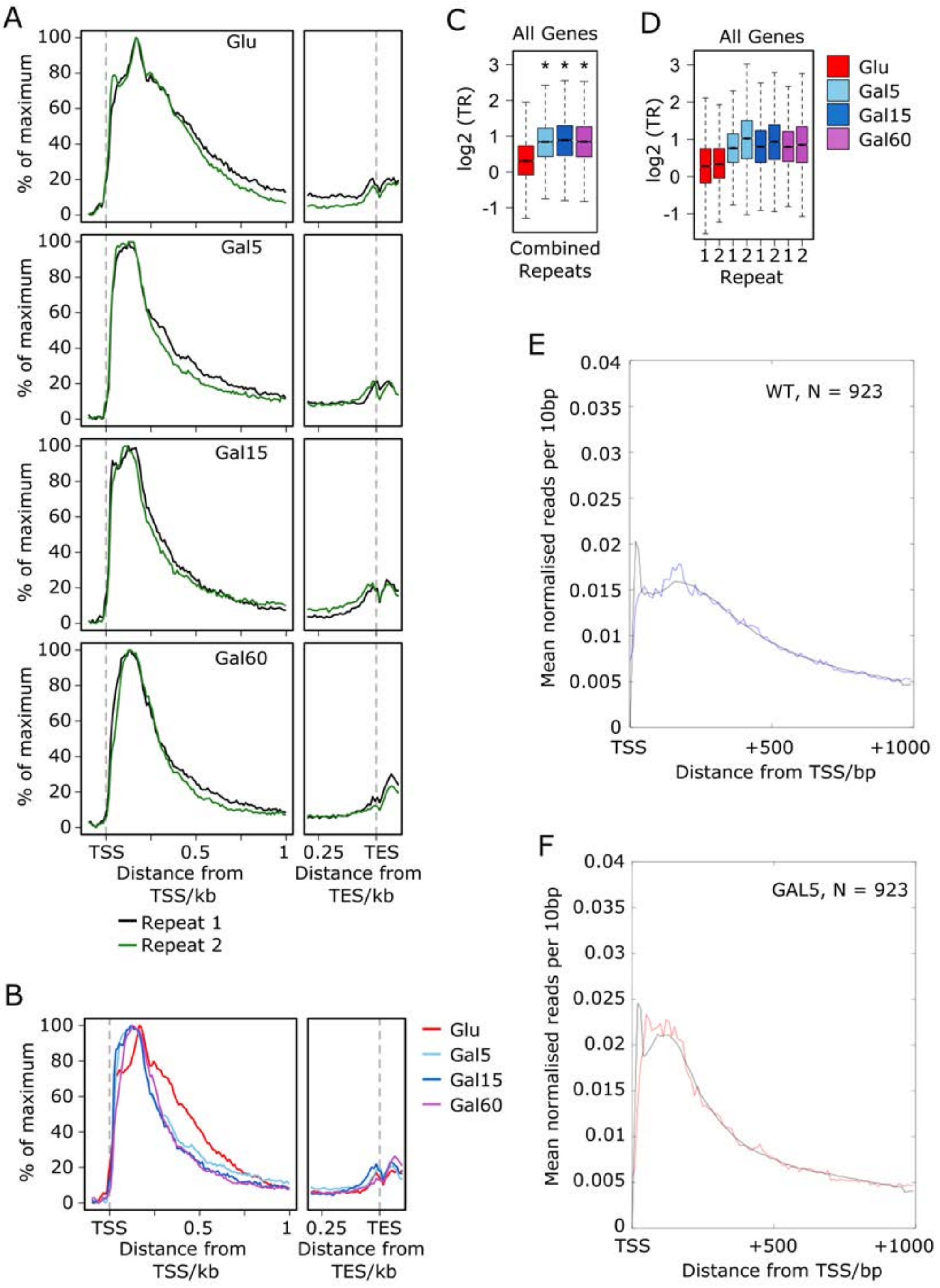

A and B) Metagene Pol2 level profiles showing the average standardized Pol2 profile for each condition for all genes for biological replicates 1 and 2 (A) and combined biological replicates (B). Profiles are calculated as detailed in Figure 6B. C and D) Boxplots showing the travelling ratios for each condition for all genes for biological replicates 1 and 2 (C) and combined biological replicates (D). Travelling ratios are calculated as detailed in Figure 6D. E and F) Dynamic parameters inferred from fitting model Metagene profiles of simulated NET-seq distributions in black after fitting to Glu profiles E) blue line or Gal5 F) redline. Note the increased promoter proximal signal in Gal5 compared to Glu.

**Table S1**

**Gene lists for each cluster and GO profiler tables for clusters 1 to 5.**

Each tab shows the gene lists for each cluster and GO classification to which each gene belongs.
